# Denoising the central autonomic network: characterizing processes jointly associated with arousal and non-neuronal physiological artefacts in dynamic fMRI analyses

**DOI:** 10.64898/2026.07.30.741873

**Authors:** Mary Miedema, Rémi Dagenais, Mohammad Torabi, S. Emad Askarinejad, Siyu Long, Georgios Mitsis

**Author notes:** Corresponding author E-mail address (M. Miedema) Affiliation address: Department of Bioengineering, McGill University, 350 McConnell Engineering Building, 3480 University Street, Montreal, QC, Canada H3A 0E9.

## Abstract

Using a multimodal dataset including fMRI, EEG-fMRI and concurrent physiological recordings, we investigated the effect of denoising systemic low frequency oscillations (sLFOs) on the characterization of dynamic signatures of central autonomic regulation and their relation to ongoing physiological states. We demonstrated that the time-frequency profiles of couplings between BOLD time series and cardiac and respiratory processes were statistically comparable between regions of the brain associated with autonomic function and non-autonomic motor regions, suggesting that these couplings largely do not reflect neuronal activation related to autonomic activity. We further showed that model-based (via physiological response functions) and data-driven (via CompCor nuisance regressors extracted from cerebrospinal fluid) methods of sLFO denoising had a statistically similar effect on these frequency profiles. We novelly applied co-activation pattern analysis to assess state dynamics of autonomic-associated regions of the brain, finding that such brain states interrelate decreases in vigilance and increases in heart rate, respiratory flow, and head motion, thus providing evidence for global arousal processes affecting BOLD signal in autonomic-associated regions. Lastly, we modelled sliding-window dynamic functional connectivity within the central autonomic network (CAN) as modulated by heart rate variability, showing significant differences in model outcomes linked to each denoising pipeline. With these findings, we provide a comprehensive discussion of the implications for the application of denoising techniques to the CAN and comment on the origins of dynamic components of the BOLD signal linked to autonomic regulation, highlighting the global role played by arousal.

## INTRODUCTION

The central autonomic network (CAN) is a set of brain regions which are implicated in autonomic nervous system function (Benarroch, 1993; Critchley et al., 2000; Sklerov et al., 2019) both in the resting state and during cognitive, emotional, or sensorimotor tasks which elicit an autonomic response (Beissner et al., 2013). The CAN reflects bidirectional neuronal processing of both efferent and afferent visceral signals relayed through key autonomic nuclei in the brainstem including the periaqueductal gray, parabrachial nuclei, and the medullary nucleus tractus solitarius, and projected to other subcortical and cortical brain structures, namely the hippocampus, thalamus, hypothalamus, amygdala, ventromedial prefrontal cortex, anterior cingulate, and insula (Beissner et al., 2013; Benarroch, 1993; Ferraro et al., 2022; Thayer et al., 2012). Sometimes implicitly, studies of the CAN have distinguished it from other autonomic-related networks such as the salience (Menon and Uddin, 2010; Sturm et al., 2018), allostatic-interoceptive (Kleckner et al., 2017; Zhang et al., 2025), and respiratory control (Krohn et al., 2023; Pattinson et al., 2009) networks by a focus on cardiovascular regulation, typically leveraging measures of heart rate variability (HRV) to identify the neural correlates of sympathetic and parasympathetic autonomic activity (Napadow et al., 2008; Valenza et al., 2024).

Using functional magnetic resonance imaging (fMRI), various abnormalities in the CAN have been observed in clinical populations experiencing autonomic dysfunction, from disruptions in connectivity in clusters of the CAN associated with sympathetic control (de Lima Xavier et al., 2025; McIntosh et al., 2021; Schultz et al., 2022; Templin et al., 2019) to increased connectivity between autonomic and brain regions involved in emotional and motor responses (Monroe et al., 2020; Thome et al., 2016; Tumati et al., 2021). The relation of resting-state connectivity to physiological autonomic measures has also been linked to emotional or homeostatic dysregulation; for instance, CAN connectivity was found to be more weakly coupled to resting HRV in participants with posttraumatic stress disorder (Thome et al., 2016) and the entropy of CAN connectivity was found to be less correlated to heart rate entropy in patients with anorexia nervosa (de la Cruz et al., 2023). Moreover, even within healthy individuals, there is convincing evidence that natural changes in physiological state affect both the strength of connectivity within the CAN and the CAN’s interactions with other brain regions at rest. It is well known that the parasympathetic and sympathetic axes of the CAN exhibit different connectivity profiles (Beissner et al., 2013; Valenza et al., 2024), and parasympathetic and sympathetic measures of HRV have been shown to differentially covary with dynamic changes in resting-state functional connectivity between regions of the CAN (Chang et al., 2013). It has previously been shown that a biofeedback task decreasing the participants’ resting heart rate (HR) and increasing their HRV also increased connectivity between the ventromedial prefrontal cortex and the amygdala (Chang et al., 2013), middle cingulate cortex, and anterior insula (Schumann et al., 2021a). Other induced changes in physiological state such as sleep deprivation (Krause et al., 2023) and stress (Huber et al., 2025; Lamotte et al., 2021) have likewise been shown to trigger functional changes within the CAN. Recently, spontaneous HRV fluctuations were shown to correspond to changes in CAN connectivity in two consecutive resting-state fMRI scans (Rominger et al., 2026), with higher HRV linked to stronger connectivity between regions of the CAN including the brainstem, insula, and anterior cingulate cortex. The relation of such intra- and inter-individual differences in sympathetic and parasympathetic outflow to CAN connectivity thus implicate fMRI measurements of the CAN as a potential biomarker.

However, studies of the CAN have rarely investigated dynamic changes in physiological state which may occur within the duration of a scan, despite well-known shifts in arousal, anxiety, and wakefulness which healthy participants frequently experience during resting-state (Gonzalez-Castillo et al., 2021; Lueken et al., 2012; Tagliazucchi and Laufs, 2014). For instance, (Chang et al., 2013) showed that HRV was positively coupled to dynamic functional connectivity (dFC) values between the dorsal anterior cingulate cortex and amygdala with other regions of the CAN and that these coupling patterns varied for different measures of HRV, possibly reflecting the influence of different sympathetic and parasympathetic processes. Several more recent studies have likewise shown that resting-state connectivity between key regions of the CAN is linearly related to changes in HR and HRV, but defining the precise nature of this relationship remains unclear, and partially attributable to the wide variety of different approaches taken to assess connectivity. While (Schumann et al., 2021a) reported that HRV was positively correlated with dFC between the ventromedial prefrontal cortex and the ventrolateral prefrontal cortex, anterior insula, and middle cingulate cortex, (Ma et al., 2024) demonstrated a negative relationship between HRV and effective connectivity between the amygdala, ventrolateral prefrontal cortex, and the anterior cingulate cortex. Additionally, (Goffi et al., 2024) identified more than 75 triplets of CAN regions whose dFC was significantly coupled with different measures of HRV. Resting-state changes in HR and FC have also been shown to exhibit a nonlinear coupling in regions associated with but not limited to autonomic control, whereby slower HR has been associated with higher FC (de la Cruz et al., 2019). Beyond the CAN itself, (Chand et al., 2020) showed that dFC between the salience, central executive, and default mode networks – which share anatomical regions and functional associations with the CAN – varies with HRV during the resting-state.

In the broader context of whole-brain fMRI, time-varying changes in physiological states have also been linked to dynamic modulation of canonical resting-state networks not specifically associated with autonomic control (Nikolaou et al., 2016). The nature of such changes in dFC is subject to ongoing interpretation (van den Brink et al., 2019) as, alongside neuronal contributions to the BOLD signal which arise through neurovascular coupling, physiological processes are well known to contribute to BOLD fluctuations through diverse non-neuronal pathways (Das et al., 2020). A recent review of methodological challenges in dFC (Laumann et al., 2024) divided this into two pathways of influence: arousal, associated to neural variability in dFC, and physiology, associated with non-neural variability in dFC; however, such a clear delineation is in practice difficult to establish, since changes in breathing patterns and heart rate frequently accompany changes in arousal. Although many data-driven and model-based strategies to remove non-neuronal physiological noise from fMRI data exist (Caballero-Gaudes and Reynolds, 2017), the extent to which denoising removes neuronal activity alongside non-neuronal physiological artefacts is difficult to assess (Bright and Murphy, 2015).

Typically, physiological denoising models remove nuisance regressors assumed to reflect physiological noise from fMRI data, leaving the “cleaned” residuals which contain BOLD fluctuations presumed to reflect neuronal processes for use in subsequent analyses (Bright et al., 2017). Such noise terms may be calculated from artefact models using concurrent physiological measurements or derived directly from fMRI data (Caballero-Gaudes and Reynolds, 2017). The use of RETROICOR nuisance regressors requires measurements of the cardiac and respiratory cycle to generate a Fourier expansion of cardiac and respiratory phase (Glover et al., 2000), which model the effect of periodic artefacts caused by physiological motion, including respiratory-induced changes to the magnetic field (Raj et al., 2001) and local cardiac pulsatility effects (Dagli et al., 1999). In contrast, systemic low frequency oscillations (sLFOs) are associated with the BOLD fMRI effects of fluctuations in systemic physiological quantities (e.g. HR and breathing) (Tong et al., 2019, 2013), with the identification of separate nonneuronal and neuronal vascular responses to these processes being non-trivial (Bolt et al., 2025; Raut et al., 2021). There exist multiple methods of denoising which remove BOLD variance associated with SLFOs; two of the most common are CompCor (Behzadi et al., 2007) and the use of physiological response functions (PRFs) convolved with measures of respiratory and cardiac activity (Birn et al., 2008; Chang et al., 2009; Kassinopoulos and Mitsis, 2019a). CompCor decomposes fMRI signals in white matter and/or cerebrospinal fluid compartments of the brain to obtain data-driven nuisance regressors; in contrast, PRF denoising offers a model-based framework for mitigating the effect of ongoing physiological changes. While global signal regression provides another potential method of denoising sLFOs, it is particularly problematic when arousal is a process of interest, as the global BOLD signal reflects relevant changes in both physiological and neuronal states (Bolt et al., 2025; Xifra-Porxas et al., 2025; Yuan et al., 2025). For instance, fluctuations in the global fMRI signal with stable temporal and spatial signatures have been observed to co-vary with arousal responses observable across various peripheral and electrophysiological signals (Bolt et al., 2024; Gu et al., 2022). Global signal regression has also been suggested to introduce spurious anti-correlations in resting-state networks (Murphy et al., 2009), although similar anti-correlations have been observed when using CompCor (Chai et al., 2012). Thus, both retaining or removing physiological artefacts from BOLD should be interpreted critically with regard to uncovering or imitating neuronal signals. It is generally true that the nuisance regressors described here contribute spatially structured noise to the BOLD signal, which may inflate connectivity when not removed (Birn et al., 2014; Tong et al., 2015). In response, the role of physiological noise in whole-brain resting-state networks has been closely examined to advocate for both the existence of neuronal circuitry and physiological mechanisms underlying intrinsic brain connectivity (Chen et al., 2020; Xifra-Porxas et al., 2021). The temporal relationship between BOLD and physiological processes provides additional evidence for a mixture of physiological and neuronal contributions to BOLD fluctuations in the sLFO frequency range. Examining the coherence between BOLD signals, (Pfurtscheller et al., 2017) reported evidence for neural BOLD oscillations occurring during periods of increased HRV alongside vascular oscillations separated by phase delay. Finally, (Lee et al., 2023) suggested that coherence between BOLD networks and cardiorespiratory measures may meaningfully identify frequency-specific autonomic contributions to network dynamics.

Returning to the CAN, since its signal of interest – i.e. parasympathetic and sympathetic activations driving physiological state change – inextricably gives rise to nonneuronal noise in BOLD signals (Das et al., 2020; Iacovella and Hasson, 2011; Macey et al., 2016), how to differentiate artefactual and neuronal measurements is a question of particular relevance for the CAN’s characterization and interpretation with fMRI. Previous studies of autonomic networks in the brain have navigated the issue of applying denoising techniques to reduce nonneuronal physiological artefacts in fMRI data with substantial heterogeneity (Kandimalla et al., 2025). While the seminal investigation relating HRV to dFC in resting-state fMRI data applied RETROICOR, CompCor, and cardiac and respiratory PRF-convolved nuisance regressors (Chang et al., 2013), few other such studies have used PRFs or incorporated such extensive denoising when considering HRV a signal of interest, whether dynamically or statically characterizing the CAN. More commonly, CompCor with varying combinations of CSF and white matter components (Jennings et al., 2016; Kong et al., 2023; Krause et al., 2023; Ma et al., 2024; Monroe et al., 2020; Rominger et al., 2026) has been used for denoising, sometimes in conjunction with other techniques such as data-driven cleaning of multi-echo BOLD data (Hohenschurz-Schmidt et al., 2020) or global signal regression (GSR) (Goffi et al., 2024). Exceptionally, one examination of dynamic changes in the CAN associated with autonomic dysfunction explicitly used a lagged RVT regressor to reduce the effect of SLFOs (de la Cruz et al., 2023), along with RETROICOR regressors. However, in other cases, no explicit physiological denoising was applied before using HRV to identify the CAN (Thome et al., 2022, 2016; Valenza et al., 2024).

Despite this wide variation in denoising approaches, it is rare for autonomic-focused fMRI studies to explicitly compare the effects of different denoising models (Miedema et al., 2025), and to the authors’ knowledge at the time of writing, no published investigation of the dynamic behaviour of the CAN has reported findings replicated across or compared between multiple denoising pipelines. It should be further noted that a recent whole-brain fMRI study reported heterogenous effects of CompCor and PRF-based denoising pipelines on FC strength and reproducibility for regions of the brain associated with arousal and autonomic control, as well as significant differences between these pipelines in terms of their sensitivity to FC changes associated with vigilance (Pourmotabbed et al., 2025). Since changes in parasympathetic and sympathetic processes exert a dynamic influence on BOLD fluctuations through both physiological artefact and neuronal processes within the CAN itself, examining the effect of denoising on fMRI measurements in autonomic regions is crucial to reproducibly characterize the time-varying behaviour of the CAN.

Thus, the present work aims to establish and interrogate the dynamic signatures of autonomic processing in the central nervous system by directly characterizing the effects of fMRI denoising associated with cardiac and respiratory ‘nuisance’ processes. Using a unique longitudinal fMRI dataset with simultaneously acquired EEG, respiratory, and cardiac data, we apply in parallel three methods of assessing dFC and physiological interactions with BOLD fluctuations in brainstem nuclei and regions of the brain associated with the central autonomic network. First, we characterize the time-varying coherence between respiratory flow (RF) and HR variations and BOLD time courses in autonomic-associated brain regions. Second, we leverage co-activation pattern analysis to associate brain states within the CAN with changes in vigilance, motion, RF, and HR. Third, we model the linear relationship between sliding-window dFC and measures of heart-rate variability and characterize its reproducibility across scans and physiological task conditions. In each investigation, we assess the effect of physiological denoising on these dynamic characterizations of autonomic interactions with the brain, demonstrating the difference made by including terms to reduce sLFO effects on BOLD fMRI data. We differentiate the use of PRF and CompCor denoising and show that sLFO-targeted denoising changes but does not eliminate signatures of autonomic outflow in CAN regions.

## METHODS

### Data acquisition

The dataset used for the present investigation was previously described in (Miedema et al., 2025). This study received approval from the McGill University Health Centre Research Ethics Board (ethics number 21-11-025; approved 17/11/2021) and written informed consent was obtained from all subjects.

Briefly, healthy subjects participated in two sessions of MRI acquisition using a 3T Siemens Prisma scanner and a 64-channel head coil. During each session, structural and functional scans were acquired along with field maps. Functional scans for the first session consisted of an eyes-open rest condition (5:09 min/300 volumes) followed by a fast-paced breathing task with interspersed breath-holds (6:52 min/400 volumes) and a passive cold pressor task (13:23 min/780 volumes). In the second session, the rest condition was repeated in a longer scan (10:18 min/600 volumes), followed by the same breathing task. For more details of each task condition, see Table 1 in (Miedema et al., 2025).

Structural and functional scans were acquired with a T1-weighted MPRAGE sequence (TE = 2.49 ms; TR = 2200 ms; flip angle = 8°; FOV = 220 mm; 0.7 mm isotropic resolution) and T2*-weighted, gradient echo planar image sequence (TE = 30 ms; TR = 1.03 s; flip angle = 46°; FOV = 220 mm; 2 mm isotropic resolution, multiband factor = 8), respectively. During functional scans, physiological data were acquired at 250 Hz (session 1) and 2000 Hz (session 2) using a Biopac MP150 acquisition module (Biopac Systems Inc, California) with an infrared sensor (Biopac, TSD200-MRI) for photoplethysmography (PPG) placed on the left (session 1) and right (session 2) thumb, as well as a respiratory effort transducer (Biopac, TSD201) placed around the abdomen. The second session included the acquisition of EEG data simultaneously during the MRI acquisition using a 128-channel geodesic sensor net and MR-compatible EEG amplifier (HydroCel GSN 230MR and Net Amps 400, EGI Inc) system sampled continuously at 1000 Hz.

Twenty-five subjects for whom complete and good quality MRI and physiological data were acquired – as assessed using in-house scripts along with MRI-QC (Esteban et al., 2017) – are included in the present analysis (mean age= 24.1±3.7 years; 13 female).

### Pre-processing

#### Physiological data pre-processing

The pre-processing of the PPG and respiratory data was performed using the pipeline previously described in (Miedema et al., 2025). To summarize, an in-house MATLAB script (available on GitHub) was used to filter the data (0.3-10 Hz and 0.01-5 Hz for PPG and respiratory data, respectively), identify peaks, correct artefact based on visual inspection and outlier removal, and calculate the following cardiac and respiratory metrics. Instantaneous heart rate (HR) was calculated as the reciprocal of the time between neighboring PPG peaks and linearly interpolated to 10 Hz. Respiratory flow (RF) was calculated as the square of the derivative of the smoothed respiratory signal and then corrected for belt slippages using an automated outlier detection routine. Lastly, from the cardiac and respiratory peaks, the phase of the cardiac and respiratory signals was calculated during each MRI volume acquisition for use in RETROICOR denoising (Glover et al., 2000).

#### EEG data pre-processing

A bandpass filter from 1-70 Hz and notch filter at 60 Hz were applied to EEG data from all channels at the time of acquisition. Subsequently, EEG data were corrected for the MR gradient artefact using average artefact subtraction (Allen et al., 2000) with a moving average template (10 TRs) implemented in Net Station 5.4 (EGI Inc). Further pre-processing of the EEG data was performed in MNE-Python (Gramfort et al., 2013; Larson et al., 2024), except where otherwise noted.

To initially correct for the ballistocardiogram (BCG) artefact, the left and right lowest EEG channels were differenced to construct a simulated ECG channel (Iannotti et al., 2015), which was adjusted in periods of low reliability using cardiac events identified using the simultaneously acquired PPG data. The simulated ECG channel was then exported along with the EEG data for use with the FMRIB plug-in for EEGLAB, provided by the University of Oxford Centre for Functional MRI of the Brain (FMRIB) to perform BCG correction with the Optimal Basis Set (OBS) method using 3 PCA components (Niazy et al., 2005). In this implementation, the lag between cardiac events and OBS-extracted windows was adjusted to be zero since the simulated ECG channel reflects the timing of cardiac effects in the EEG data, rather than cardiac events themselves.

Following these initial corrections, the EEG data were further corrected for pulse, motion, and eye-movement related artefacts with the removal of manually-labelled noisy ICA components, similar to (Gold et al., 2024) and (Yuan et al., 2013). Bad channels and time segments were excluded from the ICA decomposition to prioritize artefact reduction in good-quality EEG data (Mayeli et al., 2021). First, channels were marked as bad using automated detection to identify extreme amplitudes and a lack of expected correlation with proximal channels (Appelhoff et al., 2026; Bigdely-Shamlo et al., 2015), then interpolated with MNE-Python. Then, channels highly contaminated with noise in MR gradient- or cardiac-associated frequency ranges were discarded without replacement (outliers identified as channels higher than the median plus interquartile range of root mean square (RMS) values within 2 Hz of artefact; 10-15% of channels typically dropped). Segments of EEG data highly contaminated by noise were likewise labelled and excluded from further analysis, where time points higher than median plus six times the interquartile range of RMS values were excluded from ICA calculation and epochs containing such time points were excluded from downstream fMRI analysis (<10% of epochs dropped across all scans). The EEG data were then downsampled to 250 Hz and re-referenced to the average electrode. ICA was applied using the infomax algorithm and artefactual components were identified by visual inspection. In particular, criteria for associating components with BCG artefacts were adapted from (Gallego-Rudolf et al., 2022) as follows: 1) time series were examined for rhythmic peaks near suspected heartbeat events 2) power spectra were inspected for peaks at cardiac-related frequencies 3) topography were inspected for L-R or A-P polarity inversion, with ICs being rejected if they met two of the above criteria. Out of the 25 subjects for whom EEG data were acquired during rest and breathing scans, EEG data during 23 rest and 20 breathing scans were retained for the following vigilance-related analyses; EEG data from one rest and 4 breathing scans were excluded from further analysis due to a large majority (>75%) of their ICA components being identified as noise. One subject’s EEG data was excluded due to file corruption.

Finally, vigilance was calculated from the cleaned EEG data within epochs corresponding to each fMRI volume. The RMS value at each frequency was calculated from each epoch’s power spectral density across all channels and normalized by the RMS across all frequencies. Vigilance was then calculated as the ratio of the mean RMS value within the alpha frequency band (7-13 Hz) to the mean RMS value within the delta and theta frequency bands (1-7 Hz) (Falahpour et al., 2018; Horovitz et al., 2007; Olbrich et al., 2009; Wong et al., 2013).

#### MRI data pre-processing

Pre-processing of all MRI data was performed analogously to (Miedema et al., 2025), with the addition of brainstem-optimized registration according to recommendations from the Brainstem Navigator toolkit (Bianciardi, 2021; Singh et al., 2021, 2020). All fMRI data pre-processing was carried out using FEAT (FMRI Expert Analysis Tool) Version 6.07 in FSL39 (FMRIB’s Software Library, www.fmrib.ox.ac.uk/fsl). Initial pre-processing steps included removal of the first 3 fMRI volumes, motion realignment, fieldmap-based distortion correction with FUGUE (Jezzard and Clare, 1999), and highpass temporal filtering with a cutoff frequency of 0.01 Hz. Using the motion realignment parameters, framewise displacement (FD) was calculated for each scan volume. The functional images were co-registered to the brain-extracted high-resolution T1-weighted images using boundary-based registration (Greve and Fischl, 2009) and region-specific smoothing with a kernel of 5 mm FWHM was applied through SUSAN (Smith and Brady, 1997).

Structural images from the first scanning session were registered to the MNI152 (Montreal Neurological Institute) atlas using FNIRT (nonlinear registration with FMRIB’s Nonlinear Image Registration Tool) and resampled to 1 mm isotropic resolution. To improve the registration accuracy of small regions within the brainstem which is often neglected in favour of cortical regions (Beissner, 2015), a nonlinear transformation optimized for brainstem registration was calculated using Advanced Normalization Tools with scripts provided by the Brainstem Navigator toolkit (v 1.0; Bianciardi, 2021). This transformation was used for extracting signal from all brainstem regions. Finally, the structural data were segmented into CSF, grey matter (GM) and white matter (WM) compartments using FAST (FMRIB’s Automated Segmentation Tool).

### Application of denoising pipelines

For each pipeline, a general linear model with a set of nuisance regressors was fit to BOLD data from each scan; the residuals of this model then were retained as denoised data for use in further analysis. A baseline set of nuisance regressors was established in the **M+RETROICOR** pipeline, comprised of constant and linear drift terms, mean-centred head motion parameters along with their derivatives, and first through third order cardiac and respiratory RETROICOR terms (Glover et al., 2000). This set of regressors was selected to primarily reduce the effects of whole-head motion and local motion associated with cyclical physiological processes, including cardiac pulsatility and breathing. Two further pipelines, **+PRF** and **+CompCor**, extended this set of nuisance regressors to include terms modelling additional physiological contributions to the BOLD signal, mainly associated with sLFOs (Tong et al., 2019).

In the **+PRF** denoising pipeline, the **M+RETROICOR** regressors were combined with two terms directly derived from cardiac and respiratory measurements during each scan. Cardiac and respiratory physiological response functions (CRF & RRF, respectively) were estimated for each scan by fitting HR and RF convolved with each PRF to the global gray matter signal as described in (Miedema et al., 2025). Two further regressors were thus added to the general linear model used for denoising: HR convolved with the CRF and RF convolved with the RRF.

In the +**CompCor** denoising pipeline, the **M+RETROICOR** regressors were combined with additional terms, namely, the first five principal components extracted from BOLD time courses in CSF voxels (Behzadi et al., 2007). Although similarly-extracted white matter components are often included in aCompCor-based denoising (Kassinopoulos and Mitsis, 2019b), due to the high prevalence of white matter within the brainstem itself and to evidence for neuronally-relevant fMRI measurements within white matter (Gore et al., 2019; Grajauskas et al., 2019; Peer et al., 2017) alongside physiological effects (Duyn et al., 2020), this pipeline was restricted to the CSF compartment to best reflect a conservative application of anatomical CompCor.

### ROI definition

The brainstem serves as a site of particular interest for investigation in this study due both to its nuclei – crucial in regulating autonomic processes and interactions with the rest of the brain (Benarroch, 2018, 1993) – and its susceptibility to physiological noise during imaging (Beissner, 2015; Brooks et al., 2013; Harvey et al., 2008). Twelve ROIs corresponding to brainstem nuclei were selected from the Brainstem Navigator toolkit (Bianciardi et al., 2016; Singh et al., 2021, 2020) corresponding to the periaqueductal gray, locus coeruleus, ventral tegmental area, dorsal raphe, parabrachial nucleus, and viscero-sensory motor nuclei complex (Supplementary Materials 1). Beyond the brainstem, nineteen cortical and subcortical autonomic-associated regions (Supplementary Materials 2) were defined using spherical ROIs centred on coordinates established as centres of sympathetic and parasympathetic regulation in a previous meta-analysis of fMRI imaging during autonomic tasks (Beissner et al., 2013). Static FC for these regions during rest and breathing tasks was previously reported in (Miedema et al., 2025). Together, these brainstem and CAN-associated regions comprise a diverse set of subcortical and cortical areas in which BOLD measurements may reflect neuronal processing coupled to changes in autonomic state, with a particular focus on cardiac regulation (Figure 1). For comparison with these autonomic-associated ROIs, ‘null’ ROIs not associated with neuronal processing related to cardiac or other autonomic regulation were chosen to more specifically reflect the effects of physiological noise contributions to the BOLD signal. Three such null ROIs were selected in the brainstem (the left and right inferior olivary nucleus) and cortex (sphere centred in left primary motor cortex) (Supplementary Materials 3).

**Figure 1:**
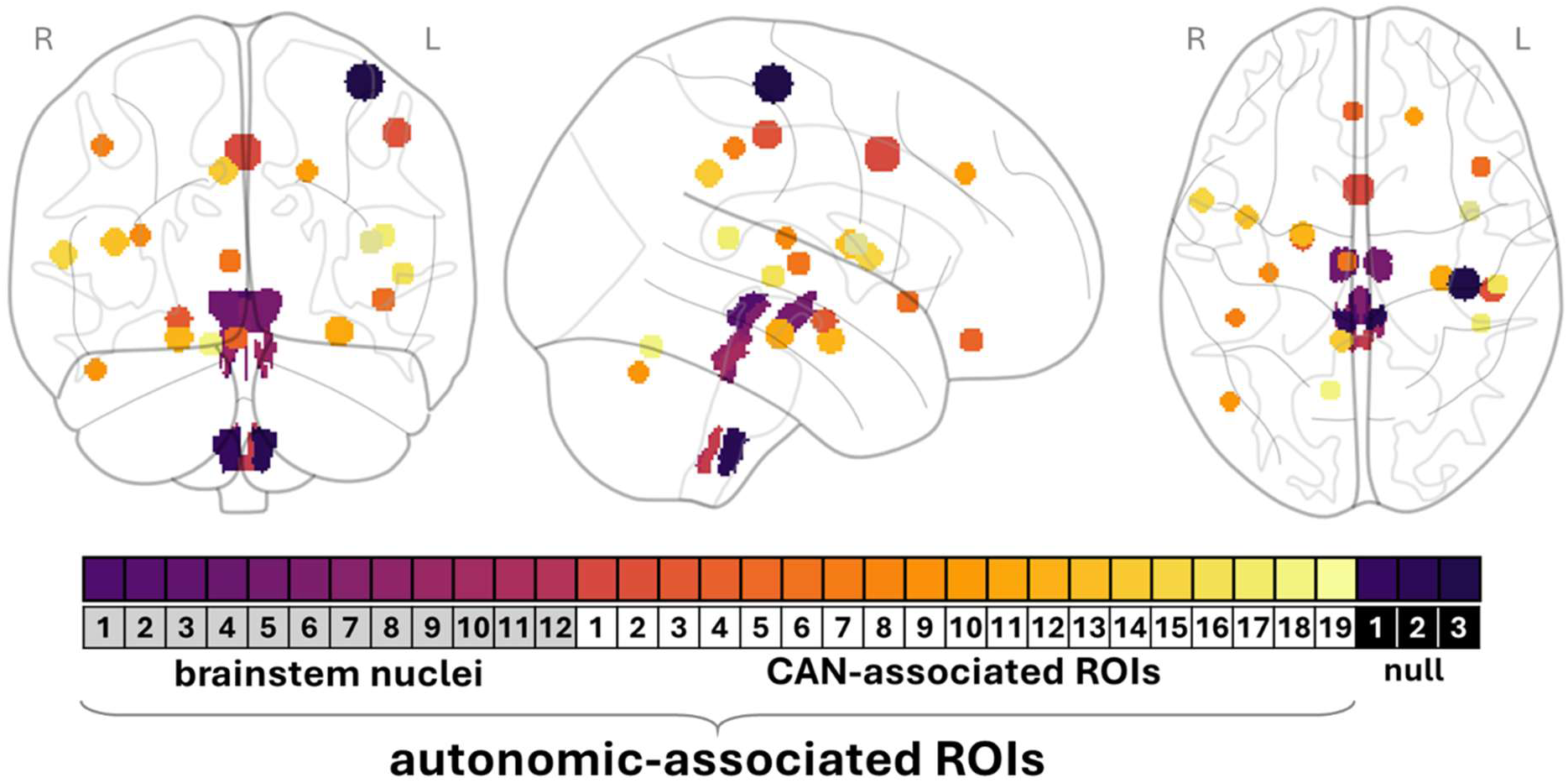
Regions-of-interest (ROIs) from which BOLD signal was extracted for the present analyses are depicted above in MNI space, along with the classification of each ROI according to their anatomical and/or functional role. The central coordinates, size, and/or labels for each ROI can be found in Supplementary Materials **1**, Supplementary Materials **2**, and Supplementary Materials **3** respectively. The CAN-associated ROIs may be further subdivided by their role in interacting with specific branches of the autonomic nervous system: ROIs 1-10 are implicated in sympathetic regulation while ROIs 11-19 are implicated in parasympathetic regulation (Beissner et al., 2013).

### Characterizations of fMRI dynamics and physiological state change

#### Wavelet transform coherence analysis

For each denoised scan, the magnitude-squared wavelet transform coherence (WTC) and the associated cross-spectrum phase was calculated between each ROI’s mean signal and concurrent HR and RF time course using analytic Morlet wavelets (Lee et al., 2023; Müller et al., 2004). The mean signal was extracted from all brainstem nuclei, all CAN-associated regions, and both sets of regions combined to examine the effect of averaging on WTC outcomes. To determine points of significant coherence between each BOLD and HR or RF time course, autoregressive models of order 1 were fit separately to each signal. Following (Chang and Glover, 2010), the estimated models and their residuals were used to generate 400 bootstrap time series pairs with the same stationary relationship, for which WTCs were calculated to obtain a null distribution for the original input signals. Significance testing was then performed with a Monte Carlo approach to threshold the magnitude-squared WTC at a 95% confidence level.

The percentage of scan volumes outside the WTC cone of influence at which significant coherence was observed was then calculated for each scan, ROI, and physiological signal. The presence of significant coherence was assessed within five frequency ranges (slow-5: 0.01 – 0.027 Hz, slow-4: 0.027 – 0.073 Hz, slow-3: 0.073 – 0.198 Hz, slow-2+: 0.198 – 0.3576 Hz) previously used to associate BOLD with electrophysiological frequency bands (Buzsáki and Draguhn, 2004; Gohel and Biswal, 2015; Zuo et al., 2010). The upper range of the slow-2+ frequency band is limited by the sampling frequency of the present study, here denoted differently than the slow-2 frequency band typically ranging from 0.198 – 0.5 Hz. Changes in the amount of significant coherence within each frequency band for each scan condition were tested for significance using a ranked Friedman test to determine whether differences between the three denoising pipelines were significant (p < 0.01, Bonferroni-corrected for multiple comparisons across frequency bands and scans). Subsequently, post-hoc Wilcoxon signed-rank tests were used to identify significant differences between each pair of denoising pipelines (p < 0.001, Bonferroni-corrected for multiple comparisons across pipeline pairs). Additionally, following (Lee et al., 2023), at each point (i.e. scan volume and frequency) for which significant coherence was identified, the phase offset of the cross-wavelet transform between HR or RF and BOLD was classified as in-phase, HR-/RF-leading, anti-phase, or BOLD-leading (0 ± π/ 4, π/2 ± π/4, -π ± π/4, and -π/2 ± π/4, respectively). The proportion of time with significant coherence at each frequency was also calculated across ROIs.

Finally, vigilance and framewise displacement were compared at time points with and without significant coherence within each frequency band during rest and breathing scans from the second session (during which simultaneous EEG data was acquired). For each ROI, physiological signal, frequency band, and scan, a ranked Friedman test was implemented to test for differences in these variables associated with coherence (p < 0.01, Bonferroni-corrected for multiple comparisons across ROIs and pipelines).

#### Co-activation pattern (CAP) analysis

To assess the relation of synchronous BOLD fluctuations across autonomic-associated regions with ongoing physiological processes, BOLD data from each scan were decomposed into sets of spontaneous co-activation patterns. Instead of following previous implementations to identify whole-brain co-activation states (Liu et al., 2018; Liu and Duyn, 2013), CAP analysis was here performed exclusively on the signals extracted from 31 autonomic-associated ROIs using the *pydfc* toolbox (Torabi, 2026). This analysis was implemented separately for each denoising pipeline, yielding a set of group-level CAP states for each pipeline and scan. For each subject’s denoised scan, these signals were z-scored and *k-*means clustered into 20 subject-level clusters. The centroids of the clusters from each subject were then combined and re-clustered to obtain group-level cluster centroids (i.e. CAP states) for each scan and denoising pipeline, where *n* - the number of group-level CAP states - was specified *a priori*. Each time point within a subject-level scan was then assigned to the most similar CAP state. This analysis was repeated to identify *n =* 2-15 group-level CAP states; for rest and breathing scans, the reproducibility of these states across sessions was assessed for each *n* by matching across sessions using correlation and then calculating average correlation across all CAP states.

The following analysis was performed only for rest and breathing scans from session 2, where simultaneous EEG data used to calculate vigilance was acquired. For each group-level CAP state, group-level distributions of HR, RF, FD, and vigilance from time points assigned to that CAP state were constructed. Since RF strongly depends on belt placement and has no inherently meaningful physiological units, it was first z-scored within each subject. For HR, FD, and vigilance, separate group-level distributions were constructed with and without subject-level z-scoring to assess CAP-state sensitivity to within-subject and group-level fluctuations in each variable. A one-way ANOVA was then used to detect differences in each variable’s distribution across all group-level CAP states.

Within each CAP application *n*, pairwise comparison testing was then used to identify CAP states associated with differences in each variable (p < 0.05, corrected for multiple comparisons across CAP states). For each such pair of states, the CAP state centroid associated with a lower mean variable was subtracted from the CAP state centroid associated with a higher mean variable, yielding a vector of region-specific differences associated with a change in that variable. The separability of changes in variables was then assessed by calculating coincidence values associated with pairs of variables within each *n*. For each variable, a state differences matrix was constructed to represent the presence and direction of changes in that variable for each pair of CAP states (Figure 7). The coincidence between two variables, representing the presence of changes in both variables associated with the same set of CAP states, was then defined as the correlation between the upper triangle of each state differences matrix.

#### Sliding window functional connectivity and heart rate variability analysis

To assess the effects of denoising and the use of different measures of HRV on assessing the modulation of dFC by autonomic processes, we fit nine linear mixed-effects (LME) models on each scan (3 pipelines, 3 HRV measures). First, dFC was calculated by correlating the BOLD time series of pairs of autonomic-associated or null ROIs within a series of sliding windows (44 TRs in duration, consecutive windows centred at each TR) with the *pydfc* toolbox. Different measures of HRV were then calculated within the same sliding windows using the simultaneous HR time series, offset in time by 3 TRs to account for the hemodynamic delay (Chang et al., 2013). First, the root mean square of successive differences (RMSSD) of heartbeat intervals (*RR*s) was calculated as follows, where *N* is the total number of interbeat intervals within each window:

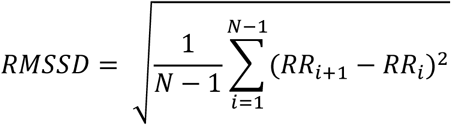

The RMSSD measure of HRV is associated with parasympathetic regulation of cardiac activity through the vagus nerve (Laborde et al., 2017) and has been suggested to be a measure of HRV more robust to breathing changes than frequency-domain measures (Thomas et al., 2019). Second, the ratio of high frequency (HF) to low frequency HRV power was calculated as an alternative measure of parasympathetic regulation. Third, the average interbeat interval (IBI) was calculated within each window. The IBI, while not a direct measure of HRV, was included in this study to investigate its nonlinear association to HRV and a marker of parasympathetic activity (Sacha, 2014) and for its relevance to previous investigations of autonomic-dFC coupling (de la Cruz et al., 2023; Sonkusare et al., 2025). Of these measures, the RMSSD was additionally log-transformed to compensate for skewed data distributions.

For each of these three variants of HRV, measures were z-scored within each subject to obtain an HRV_state_ variable calculated from each sliding window, while the mean HRV within each subject across all sliding windows was used to define the variable HRV_trait_. Additionally, within each window, FD was summed to obtain a variable reflecting motion-specific effects which was then log-transformed to account for skew and z-transformed across all subjects. Finally, an LME model with random effects for subject intercepts was fit for each pair of ROIs (i.e. dFC edge) according to the following formula:

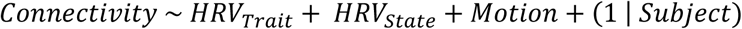

The model fit was implemented using the *fitlme* function in MATLAB R2023a (The MathWorks, Natick, USA). For statistical testing of the beta coefficients obtained for HRV_state_ in the linear mixed-effects model, surrogate HRV data with phase randomization within each subject were used to create a null *t-*distribution to assess significance across 2000 realizations. In terms of modelling the dFC patterns between the 31 autonomic-associated ROIs and 3 null ROIs selected for this study, results are reported for 561 pairwise connections without correction for multiple comparisons. This approach was chosen to maintain sensitivity of these results to differences between pipelines and scans which may be relevant for future studies interrogating specific regions within the CAN or brainstem; outcomes in the present study should therefore be evaluated carefully in relation to type one errors.

The cosine similarity of the LME beta matrices (thresholded to the same significance level) was calculated between all pairs of pipelines and scans to assess the reproducibility of model outcomes. Additionally, for each scan, pipeline, and significance level, undirected node strength (i.e. the sum of significant beta values for each ROI) was calculated using the *strengths_und* function in the Brain Connectivity Toolbox (Rubinov and Sporns, 2010). ROIs were then ranked by their node strength and the Spearman rank correlation of these rankings was calculated to compare model outcomes across scans and pipelines.

## RESULTS

### Coherence of BOLD signals with heart rate and respiratory flow

#### Effects of denoising and scan type on coherence with RF and HR

The percentage of scan volumes which exhibited significant coherence with each physiological signal is shown for all autonomic-associated regions in Figure 2. Similar results were observed for null regions (Supplementary Materials 4A); a dedicated comparison between the coherence patterns in null and autonomic-associated regions is made in the following section. Averaging the percentage of scan volumes in coherence for each frequency across all autonomic-associated ROIs following their cross-wavelet transforms with HR and RF, the M+RETROICOR pipeline resulted in the greatest such coherence within most frequency bands (Figure 2A). The percentage of each scan in coherence with RF was equivalently reduced by PRF and CompCor denoising within most frequency bands, except within the slow-4 frequency band, where CompCor denoising reduced coherence more than PRF denoising during the cold pressor and long resting state scans (Figure 3). Likewise, coherence with HR was similarly reduced by PRF and CompCor denoising within most frequency bands, although CompCor denoising reduced coherence with HR more than PRF denoising within the slow-5 and slow-4 frequency bands during the breathing and long resting state scans, respectively. Notably, during the breathing task, the distribution of HR and RF coherence was shifted towards the slow-5 and slow-4 frequency bands across all denoising pipelines. This was expected due to the strong physiological stimulus and the effects of respiratory sinus arrhythmia, which increased the proportion of scan volumes exhibiting significant coherence with both HR and RF. In the resting-state, our results showed a similar distribution in coherence with HR to that reported by (Lee et al., 2023) when using the heartbeat interval.

**Figure 2:**
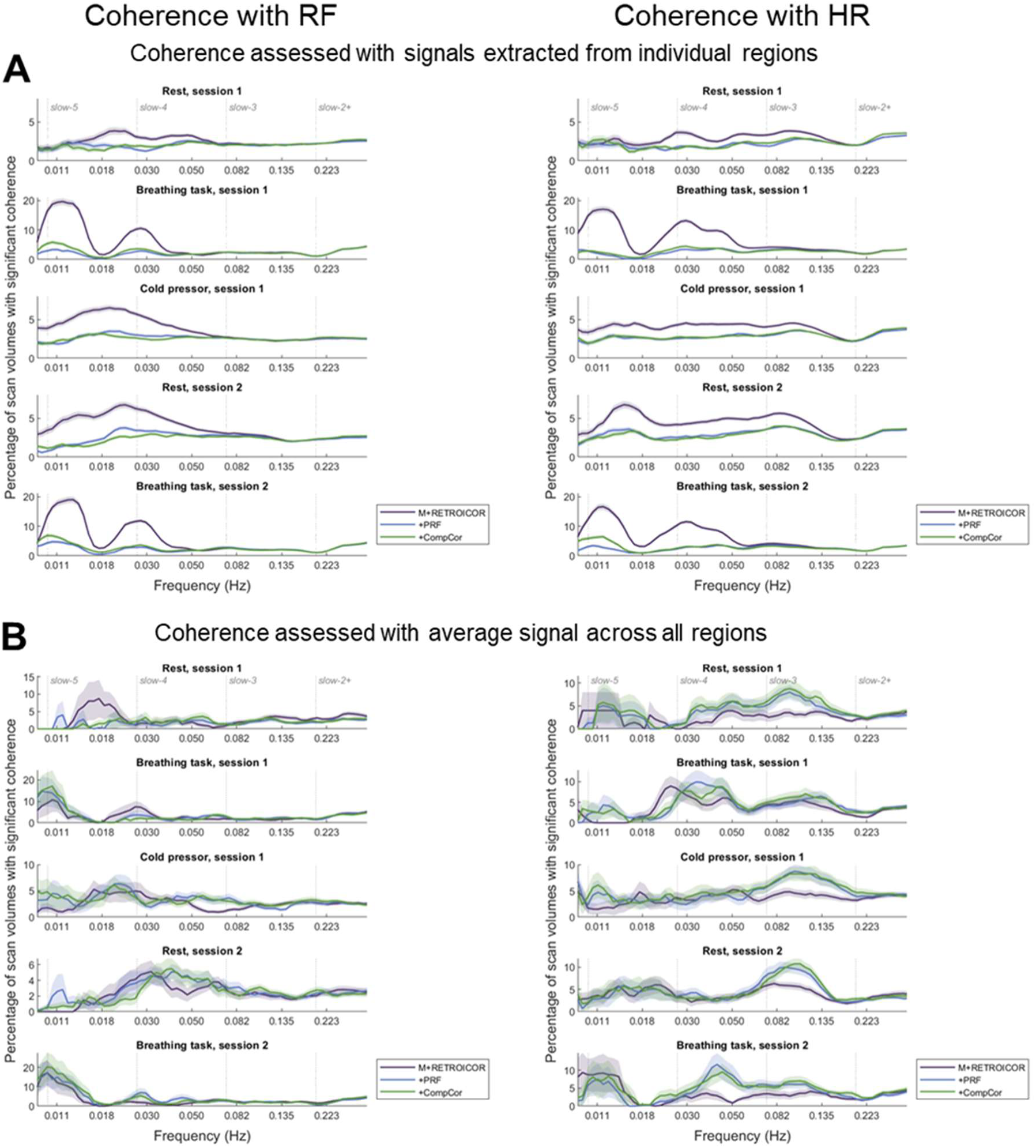
M+RETROICOR denoising results in greater coherence with HR and RF in frequencies compatible with neuronally modulated hemodynamic responses (slow-5, slow-4, and slow-3 bands) when the cross-wavelet transform is computed for individual autonomic-associated ROIs (A), but this is not the case when the BOLD signal is averaged across all autonomic-associated ROIs prior to computing coherence (B). The percentage of scan volumes which exhibited significant coherence is plotted against frequency for each scan and denoising pipeline. Shaded regions represent the standard error for each pipeline and scan, over participants and ROIs (A) or over participants only (B).

**Figure 3:**
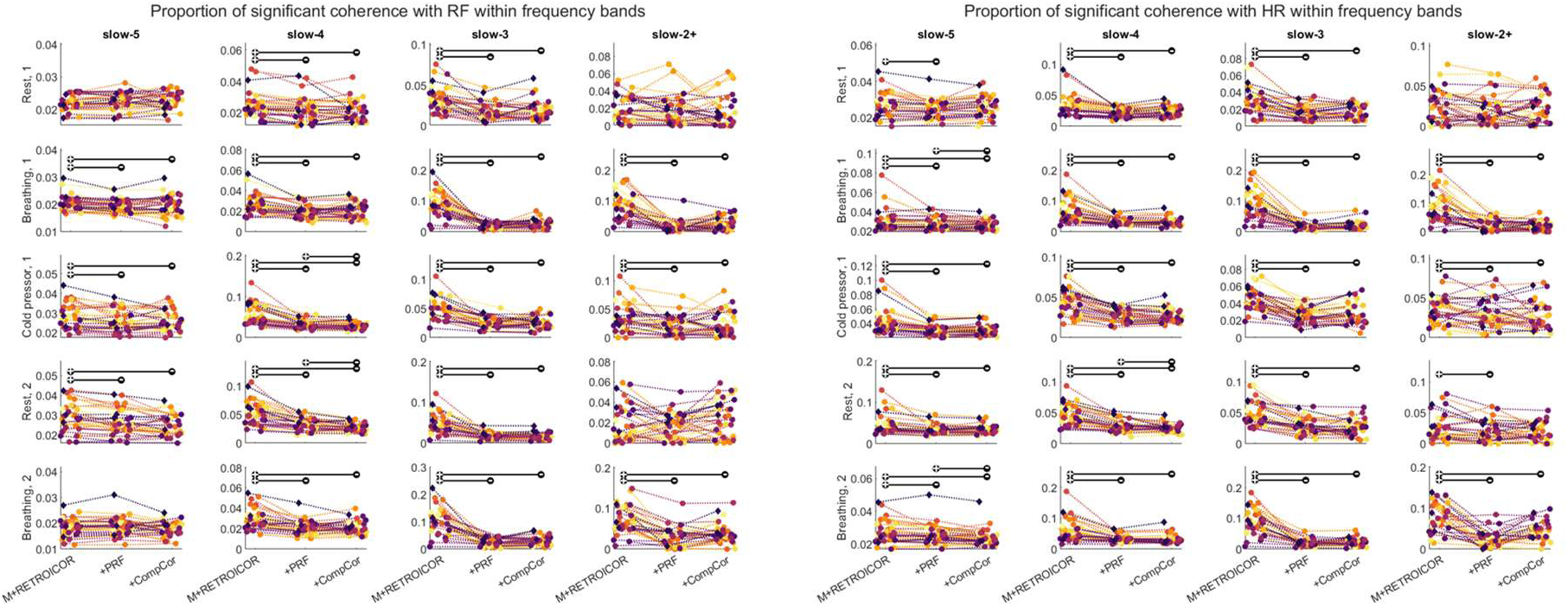
PRF and CompCor denoising significantly and similarly reduce the amount of coherence with RF and HR in neuronally-relevant frequency bands. The proportion of scan volumes with significant coherence is plotted for each ROI (marker coloring reflects ROI definitions as per Figure 1), denoising pipeline, scan, and frequency band. The effect of denoising pipeline on the proportion of scan volumes exhibiting significant coherence with each physiological signal is shown with bars indicating greater (+) and lower (-) amounts of coherence across all ROIs (p < 0.001, Bonferroni-corrected for multiple comparisons). Null ROIs (diamonds) are included for reference in this visualization, but were not included in statistical testing, which was performed only across autonomic-associated ROIs (circles).

In contrast, when signals across autonomic-associated regions were averaged prior to application of the cross-wavelet transform, the sLFO-denoising pipelines displayed a tendency to increase coherence with HR but not RF relative to the M+RETROICOR pipeline (Figure 1B). Averaging the BOLD signal across ROIs prior to assessing coherence significantly increased the amount of coherence with HR at higher frequencies, particularly for PRF and CompCor denoising during the rest and cold pressor scans. In contrast, coherence with RF was typically slightly reduced (with the exception of M+RETROICOR denoising during rest, session 1) and became more similarly distributed across denoising pipelines. These results suggest that the sLFO-denoising pipelines may remove respiratory fluctuations which are shared globally across ROIs but potentially introduce systemic cardiac fluctuations when denoising is performed on spatially distributed regions prior to averaging. Moreover, brainstem-specific effects were observed when averaging was performed within specific subsets of autonomic-associated regions (Supplementary Materials 4B).

Although the sLFO pipelines had similar effects on amount of coherence observed within several frequency bands, the times at which significant coherence was identified within each scan differed with each denoising pipeline except at low (slow-5) frequencies (Supplementary Materials 5). Dice overlap scores and Euclidean distance were used to compare whether pipelines resulted in coherence that was identified at the same times and frequencies within each scan and ROI. Although the Euclidean distance between the coherence patterns for different pipelines was similarly distributed across frequency bands (Supplementary Materials 5B), Dice overlap scores neared zero in the slow-4 to slow-2+ frequency bands (Supplementary Materials 5A), highlighting the strong effect of denoising on coherence significance testing. Across all frequency bands, Dice overlap scores were higher during the resting and cold pressor scans than during the breathing task, indicating that differences between denoising pipelines were exacerbated by the presence of strong physiological artefacts (Supplementary Materials 5C).

When significant coherence was observed, the phase offsets between the BOLD signal in CAN-associated regions and RF/HR were modulated by tasks and denoising within neuronally-relevant frequency bands (Supplementary Materials 6). As expected, since PRF denoising regresses a convolved (i.e. temporally delayed) RF and HR from the BOLD signal; a corresponding reduction in the proportion of RF- or HR-leading coherence was observed across tasks in the slow-3 band, but was frequently accompanied by an increase in HR-leading coherence in the slow-5 or slow-4 bands (peak ∼ 0.03 Hz). This same effect was observed to a lesser extent overall in the case of CompCor denoising, whereby the most prominent effects were observed during the breathing scans. During these scans, M+RETROICOR coherence was highly in-phase within the slow-4 band, while for frequencies lower than 0.02 Hz anti-phase and BOLD-leading coherence were mostly observed. In general, CompCor denoising reduced the differences in phase proportions between rest and breathing tasks more than PRF denoising, for which peaks in anti-phase coherence could still be observed in the slow-5 band during breathing tasks.

#### Specificity of coherence to autonomic-associated regions

To clarify the interpretation of coherence with physiological signals, especially HR, as indicative of neuronal or non-neuronal processes measured by BOLD, the coherence of autonomic-associated regions was compared to that of null regions within each frequency band (Figure 4). Separate comparisons were made for brainstem nuclei and for other CAN-associated cortical and subcortical regions in light of the brainstem’s particular susceptibility to physiological noise (Harvey et al., 2008).

**Figure 4:**
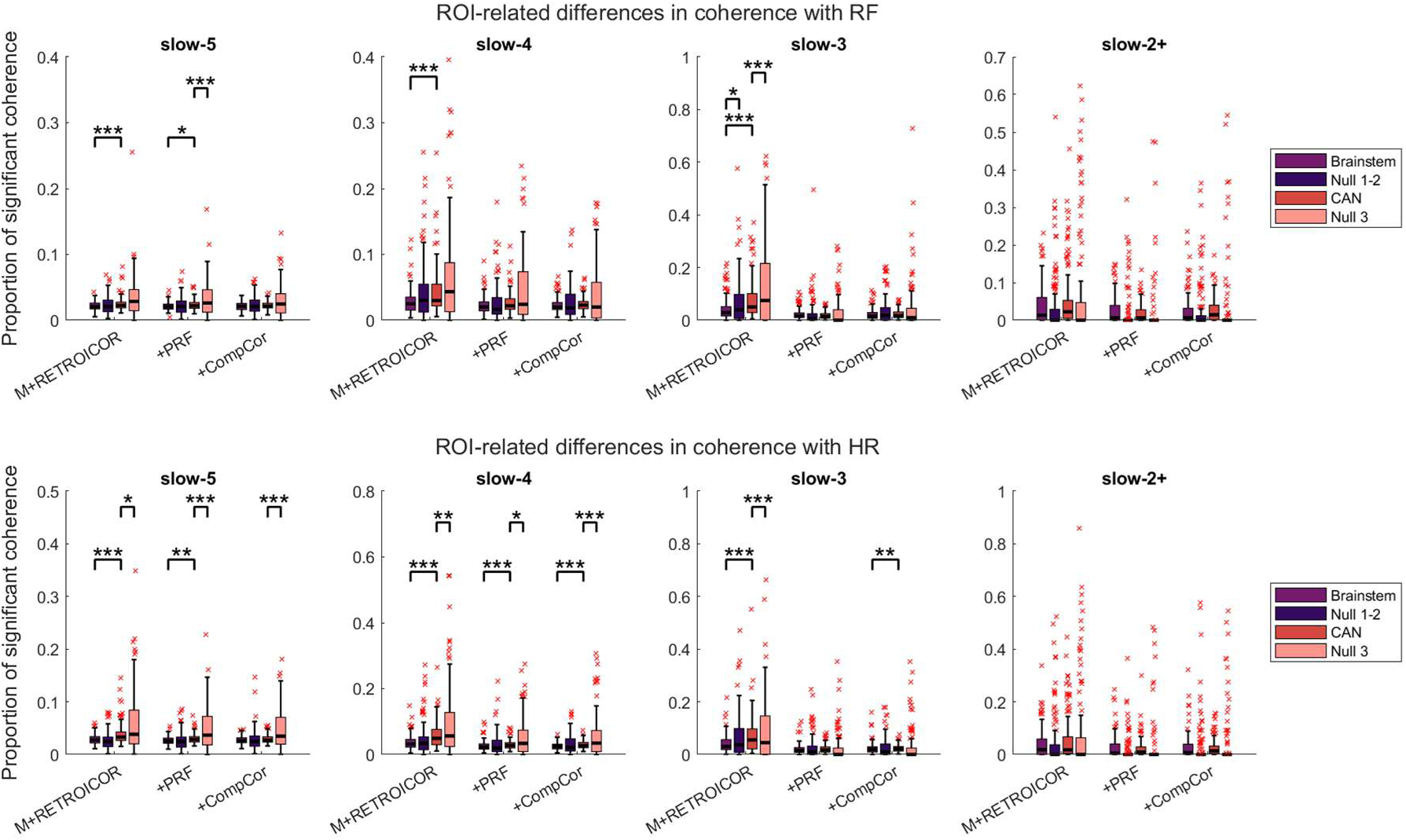
**Autonomic-associated regions do not generally show higher coherence with physiological signals than matched null regions**. The distributions of the average proportion of significant coherence within each frequency band is plotted across subjects and scans for each region type. Statistically significant differences are shown on each plot, as assessed with a ranked Friedman test within each pipeline to test for differences across regions (p < 0.01, Bonferroni-corrected for multiple comparisons across frequency bands and pipelines) followed by post-hoc Wilcoxon signed-rank tests (p < 0.05 = *, < 0.01 = **, < 0.001 = ***).

For RF, similar amounts of coherence were found across ROIs of all types for most pipelines and frequency bands, although for M+RETROICOR in the slow-5, slow-4, and slow-3 bands CAN ROIs were found to exhibit greater coherence than brainstem ROIs (p < 0.001). In the slow-3 band alone, both null ROI types were found to have greater coherence with RF than their brainstem and CAN counterparts for the M+RETROICOR pipeline. Additionally, in the slow-5 band for the +PRF pipeline, CAN ROIs were found to have greater coherence than brainstem ROIs (p < 0.05) and the null motor ROI was found to have greater coherence than CAN ROIs (p < 0.001).

For HR, the null motor ROI was found to have greater coherence than CAN ROIs in the slow-5 and slow-4 bands across all pipelines. Likewise, in the slow-5 and slow-4 bands, CAN ROIs were typically found to have greater coherence than brainstem ROIs. This CAN versus brainstem trend persisted into the slow-3 band for the M+RETROICOR and +CompCor pipelines; in this band and for the M+RETROICOR pipeline alone, CAN ROIs exhibited greater coherence with HR than the null motor ROI.

Relating coherence with heart rate and respiratory flow to changes in vigilance and motion To assess how BOLD signal coherence with cardiorespiratory regulation should be interpreted in relation to arousal and motion, differences in the distributions of vigilance and FD stratified by coherence with HR and RF were assessed for each ROI independently. The results that follow arise from data during session 2 only, since no EEG data were acquired during session 1. The typical correlation between HR, RF, vigilance, and FD time series during session 2 was low, with an average value < 0.2 across scans and subjects (Supplementary Materials 7).

Significant changes in vigilance occurring during coherence with RF and HR were observed for the majority of ROIs, with some general trends observable within frequency bands and preprocessing pipelines (Figure 5A&B). First, for both RF and HR, coherence in lower frequency bands (slow-5 to slow-3) was more associated with reduced vigilance, while coherence at higher frequencies (slow-2+) was more associated with increased vigilance. Also, the magnitude of the observed differences in vigilance increased for some regions at higher frequencies. Second, at lower frequencies (slow-5 and slow-4), CompCor denoising tended to reduce the number of significant differences in vigilance observed relative to M+RETROICOR and PRF denoising. Third, coherence with null and mean ROIs (where signals were averaged across multiple ROIs prior to coherence calculation, c.f. Figure 2B) also often corresponded to significant changes in vigilance. Overall, the finding that coherence with physiological signals below 0.198 Hz was associated with lower vigilance is consistent with a previous study which demonstrated stronger coupling between BOLD signals with heart rate and respiration during sliding windows with lower vigilance (Gold et al., 2024).

**Figure 5:**
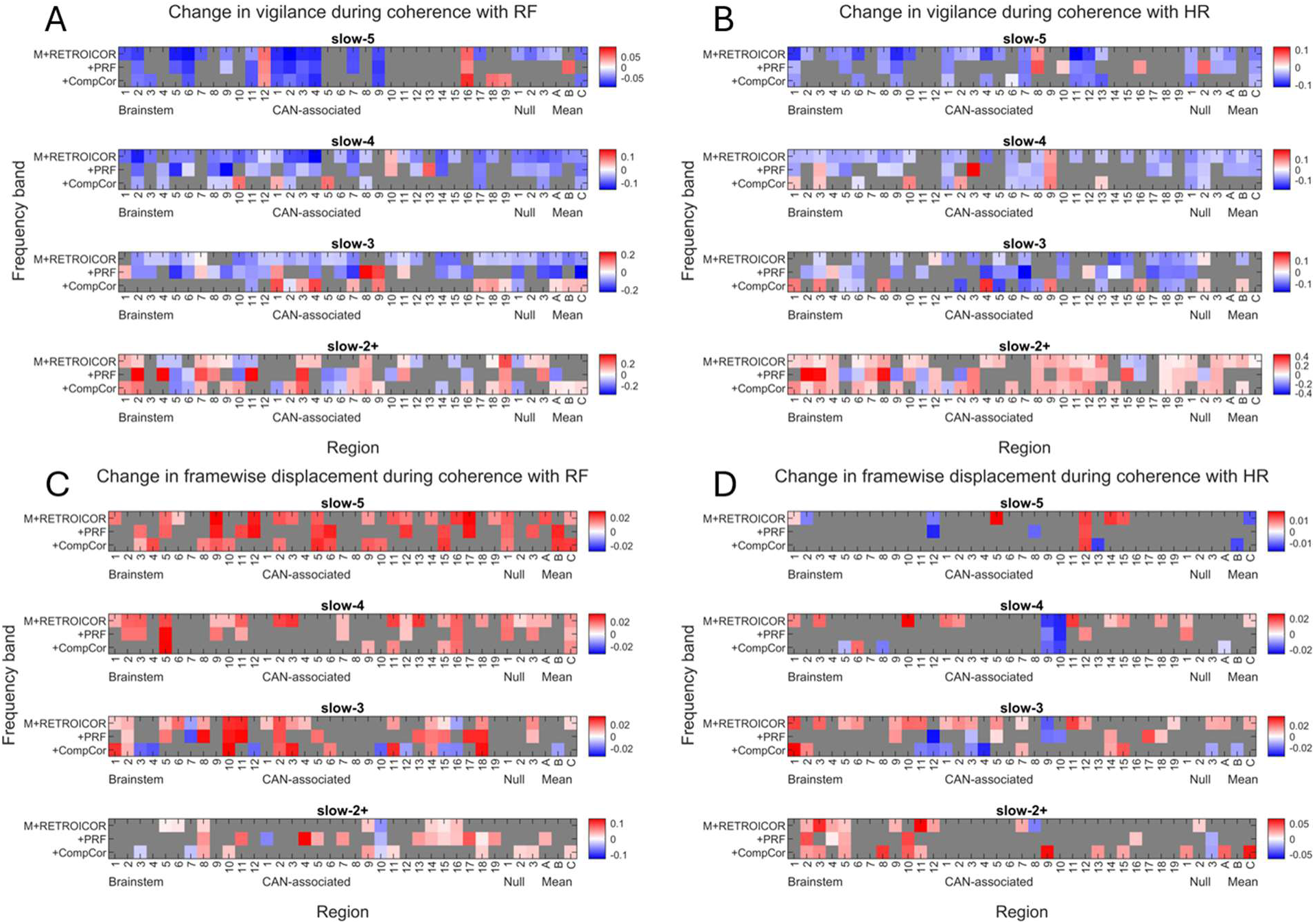
Fluctuations in vigilance and FD are linked to the presence of coherence with RF and HR within individual ROIs. Magnitude of significant differences in vigilance (A&B) and FD (C&D) distributions during periods of significant coherence with RF (A&C) and HR (B&D) as assessed with a Wilcoxon rank sum test (p < 0.01, corrected for multiple comparisons within each frequency band). Labels of regions correspond to their designations in Tables 2-4; mean region labels denote coherence with signals which were first averaged over all autonomic-associated regions (Mean-A; Tables 2-3), brainstem nuclei (Mean-B; Table 2), and CAN-associated regions (Mean-C; Table 3).

In contrast, significant changes in FD occurring during coherence with HR were observed for fewer ROIs than for coherence with RF (Figure 5C&D)., consistent with the stronger motion artefacts induced by breathing. This was particularly true for lower frequency bands (slow-5 and slow-4), in which coherence with HR in only a few ROIs was linked to significant changes in FD (Figure 5D). At higher frequencies (slow-2+), the magnitude of the observed differences in FD also increased for coherence with both HR and RF. Unlike vigilance, changes in FD were largely positive during coherence with both HR and RF.

For both differences in vigilance and FD, the effects of denoising within a given frequency band and ROI were often variable. In numerous instances, the application of a different denoising pipeline changed the strength or even the direction of changes in vigilance or FD during periods of significant coherence with physiological fluctuations. It is notable that while the addition of sLFO denoising reduced the presence of significant coherence-associated differences in vigilance and FD within certain ROIs and frequency bands, the converse was also frequently observed: PRF or CompCor denoising sometimes resulted in significant coherence-associated differences not present in the M+RETROICOR pipeline.

Despite the difficulty of generalizing from such variable results, close examination of cases when the three pipelines obtained similar coherence-associated differences within a given ROI could provide insight into possible processes which drive coherence with physiological signals. For instance, although coherence with RF in the slow-5 band was associated with lower vigilance, higher vigilance can be seen during coherence with RF in the right viscero-sensory-motor (VSM) nuclei complex (Brainstem ROI 12) and temporal gyrus (CAN-associated ROI 16). This positive association with vigilance in the right VSM and negative association with vigilance in the left VSM (Brainstem ROI 11) suggests a meaningful neuronal lateralization of coherence with RF specific to this brainstem region, since this effect is not present in other nearby brainstem nuclei which should be similarly susceptible to lateralized physiological artefact. Previous fMRI studies in rodents have demonstrated a right-sided subregion in the nucleus of the solitary tract activated by hypoxia (MacMillan et al., 2024; Mahmoud et al., 2016); since the VSM contains this region, the neural response to respiratory demand of fast paced breathing and breath-holds during the breathing task scan may therefore explain this effect. The BOLD signal in the temporal gyrus, on the other hand, was previously identified as negatively correlated with both EEG alpha power and respiration during eyes-closed rest (Yuan et al., 2013) and is further implicated in responding to vigilance changes through its role in the ventral attention network (Das et al., 2026). Increased vigilance during RF coherence with the temporal gyrus therefore suggests that coherence with RF in this region may reflect at least in part a neuronal contribution to the BOLD signal.

Likewise, some ROIs were found to be highly susceptible to motion-related artefact during coherence within certain frequency bands across all denoising pipelines. It is particularly interesting that FD was lower during coherence with HR in the cerebellum and dorsolateral prefrontal cortex (CAN-associated ROIs 9 & 10) in the slow-4 frequency band; it is possible that such reductions in FD during coherence reflect ROIs especially susceptible to motion artefacts, which fail to exhibit coherence with physiological signals during periods of elevated movement. In contrast, ROIs exhibiting higher FD during coherence seem more likely to track changes in global arousal processes that correspond to periods of higher motion; the broad involvement of coherence with RF across ROIs suggests a strong effect of chest and head motion while performing the breathing task.

Finally, in examining the behaviour of null ROIs and signals averaged across autonomic-associated ROIs (‘mean’ ROIs), we note that the presence of vigilance and FD-associated differences during coherence is not specific to individual autonomic-associated ROIs. In many cases, coherence with HR and RF in null ROIs (or when using the average brainstem or CAN BOLD signals) gives similar results to those discussed above, with similar variability introduced by denoising. Overall, these results suggest that coherence with HR and RF in most frequency bands and ROIs is indicative of contributions made by mechanical physiological artefacts or globally-mediated neuronal processes associated with physiological arousal, rather than region-specific neuronal activations reflecting local autonomic processing. However, as previously gestured towards, several of our results could be consistent with coherence reflecting meaningful neuronal involvement, suggesting that this technique could be used to investigate mechanisms of interaction with physiology. The magnitude of changes in vigilance associated with coherence in the slow-2+ band for several ROIs in PRF denoising also present a possible target for further study. Our results do not preclude, for instance, that the increase in vigilance observed during coherence between HR and the locus coeruleus (Brainstem ROIs 2 & 3) reflects an increased sensitivity to neural activation during shifts in arousal.

### Examining BOLD signal changes across autonomic-associated regions with co-activation pattern analyses

#### Effects of denoising on co-activation pattern sensitivity to physiological changes

For each CAP analysis (*n* states 2-15), we assessed whether pairs of CAP states could be associated with differences in each of the four measures used for quantifying ongoing physiological processes. Distributions of vigilance, FD, RF, and HR stratified by each CAP state were therefore tested for significant differences amongst all CAP states identified for each denoising pipeline (Figure 6), for rest and breathing scans acquired during session 2.

**Figure 6:**
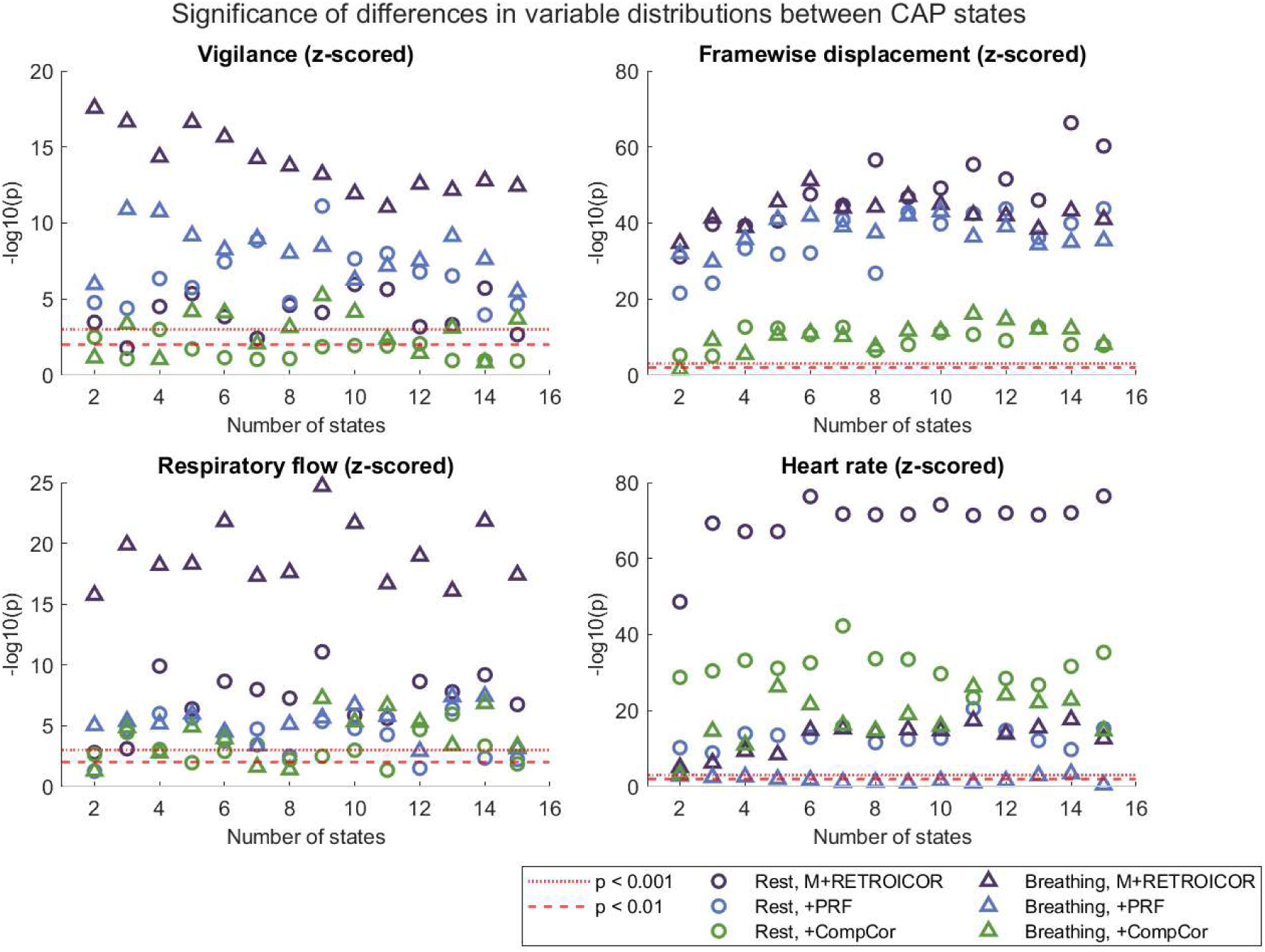
The significance of differences in vigilance, FD, RF, and HR distributions corresponding to different CAP states vary with task condition and denoising pipeline. The *p*-values from a one-way ANOVA are plotted here for each number of group-level CAP states (*n* = 2…15), for *z-*scored physiological variables’ distributions corresponding to CAP decompositions of rest and breathing scans acquired during session 2.

Differences in vigilance between CAP states were identified using the M+RETROICOR and PRF pipelines for all *n* (p < 0.05) during the rest and breathing scans. In contrast, for the CompCor pipeline, differences in vigilance could not be detected between CAP states in the resting-state for *n > 2* and were inconsistently detected during the breathing task. CompCor denoising also reduced the significance of differences in FD during rest and breathing relative to the M+RETROICOR and PRF pipelines, although for all pipelines, differences associated with motion could be identified between CAP states (p < 0.05). Differences in respiratory flow were identified for all *n* in the M+RETROICOR pipeline (p < 0.05) but sLFO denoising resulted in no significant differences in RF in several instances. Finally, differences in HR were typically significant for all pipelines in the resting-state, but during the breathing task no differences in HR could be detected for CAP states denoised with PRFs. However, differences in HR were similarly significant for PRF and CompCor denoising when inter-subject differences in HR were not accounted for (c.f. results presented without *z*-scoring in Supplementary Materials 8). The significance of differences in HR was greatly reduced by both sLFO denoising pipelines. Differences in HR were more significant during the resting state than during the breathing task for all pipelines and number of CAP states, a result replicated for n > 5 without *z*-scoring, suggesting variability in arousal was greater during rest than breathing modulation.

#### Specificity of co-activation patterns to different physiological processes

We examined the ROI-specific differences obtained from subtracting the centroid vectors of pairs of CAP states for which a significant difference was associated with vigilance, FD, RF, or HR; when multiple such pairs occurred within a given *n*, the average difference over all pairs was obtained weighted by the difference in vigilance, FD, RF, or HR distributions. The ROI-specific differences associated with a positive change in each measure is shown in Figure 7A for each pipeline and *n*. Certain ROIs seem to consistently drive the identification of CAP states associated with differences in these physiological processes: namely, the cingulate cortex, supramarginal gyrus and somatosensory cortex, and ventrolateral prefrontal cortex (CAN-associated ROIs 1, 2, 5 & 7); similar results were obtained without *z-*scoring (Supplementary Materials 9). During the breathing task, differences of greater magnitude are observed, particularly for the M+RETROICOR pipeline. In contrast, sLFO denoising tends to reduce the magnitude of variable-associated differences, particularly for the CompCor pipeline. While brainstem ROIs seem not to play a significant role in differentiating CAP states associated with physiological changes for M+RETROICOR and PRF denoising, differences in these regions are more comparable to other CAN-associated ROIs in the CompCor pipeline. One notable exception in the PRF pipeline are the differences associated with changes in RF in the resting-state, where positive BOLD fluctuations in brainstem regions seem linked to increased RF.

**Figure 7:**
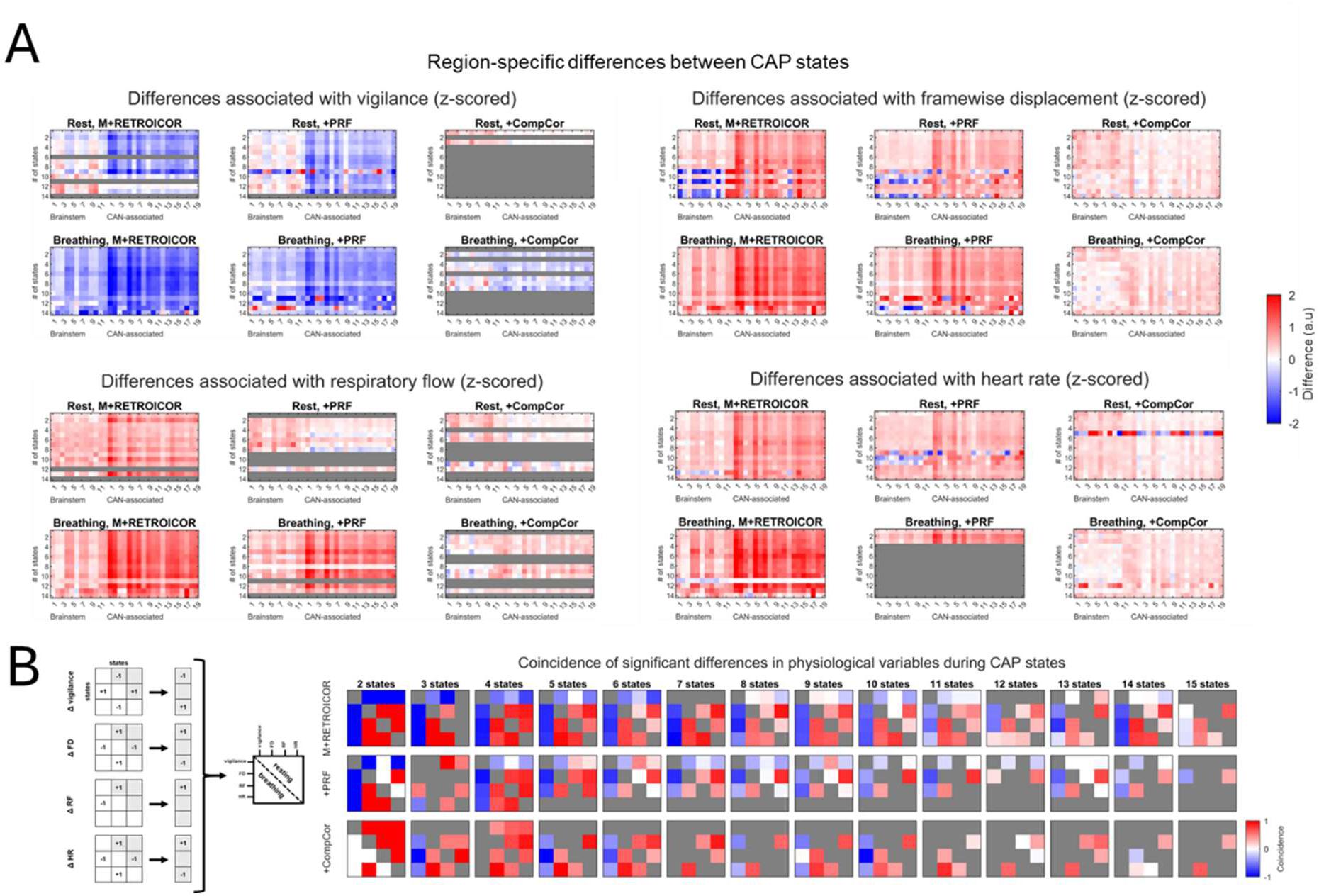
Differences in CAP state centroids primarily link fluctuations in non-brainstem regions to changes in vigilance, FD, RF, and HR. Differences between centroids are plotted to show the change in each ROI only for pairs of CAP states exhibiting a change in physiological variable (A) while coincidence in changes between each pair of physiological variables is assessed across all CAP states (B). Gray regions represent instances where no significant differences in the distribution of a variable were observed across all CAP states.

To further assess whether changes in each measure were correlated with changes in other measures, for each *n* and measure we constructed a matrix reflecting the presence of significant differences between states (Figure 7B). The correlation of the upper triangles of these matrices was then calculated to assess the coincidence of changes in each pair of measures. A coincidence value of zero indicates that a change in measure A never occurred between the same pair of CAP states for which a change in measure B occurred. In contrast, a coincidence value of one indicates that a change in measure A occurred for the same set of states in which a change in measure B occurred, and that these changes occurred in the same direction. A coincidence value of negative one indicates that changes in measures A and B occurred in the same set of states, but in opposite directions. In Figure 7, several trends emerge across pipelines, tasks, and values of *n*. First, changes in physiological variables coincide more during the breathing task than at rest, as expected due to the strong physiological couplings introduced by this task. However, changes in physiological variables frequently coincided during rest, and denoising did not usually improve the separability of changes in physiological variables. For instance, a positive coincidence between FD and HR was consistently observed at rest as well as during breathing scans; the coincidence value decreased with *n* and was most decreased in the CompCor pipeline but remained similar in the PRF and M+RETROICOR pipelines. Typically, increases in vigilance were often accompanied by decreases in motion, respiratory flow, and heart rate, although using a larger value of *n* improved the separability of changes in these measures, and the PRF pipeline likewise sometimes reduced coincidence with vigilance relative to the M+RETROICOR pipeline.

Overall, these results highlight the tendency of each denoising pipeline to remove or preserve BOLD fluctuations associated with different physiological processes. While CompCor denoising best mitigated the effects of motion, it also reduced variance in the BOLD signal linked to changes in vigilance (Figure 6). In contrast, PRF denoising did not allow the detection of CAP states which discriminated intra-subject changes in heart rate during breathing task performance, although this was not the case for absolute changes in HR (Supplementary Materials 8). Finally, higher dimensionality implementations of CAP allowed greater separability of changes in physiological variables associated with specific CAP states, which tended to coincide for lower *n* for all denoising pipelines, suggesting that greater *n* could be advantageous for isolating neural autonomic activity.

### Heart rate variability modulation of sliding window connectivity

#### Sliding window analysis of dynamic functional connectivity

The mean values and standard deviations of dFC between ROIs across all sliding windows are shown for each scan and denoising pipeline in Figure 8. In the M+RETROICOR pipeline, the mean FC across all sliding windows was higher within CAN-associated regions and null regions than within brainstem regions and for connections between brainstem and CAN-associated regions. The magnitude of this difference in mean FC varied with scan type, increasing with longer scans. Denoising tended to reduce the difference in mean FC across region types; the CompCor pipeline did so more aggressively than the PRF pipeline. However, differences in the variability of dFC across sliding windows remained across all denoising pipelines: connections within CAN-associated regions and null regions were significantly more variable than connections with brainstem regions for all scan types.

**Figure 8:**
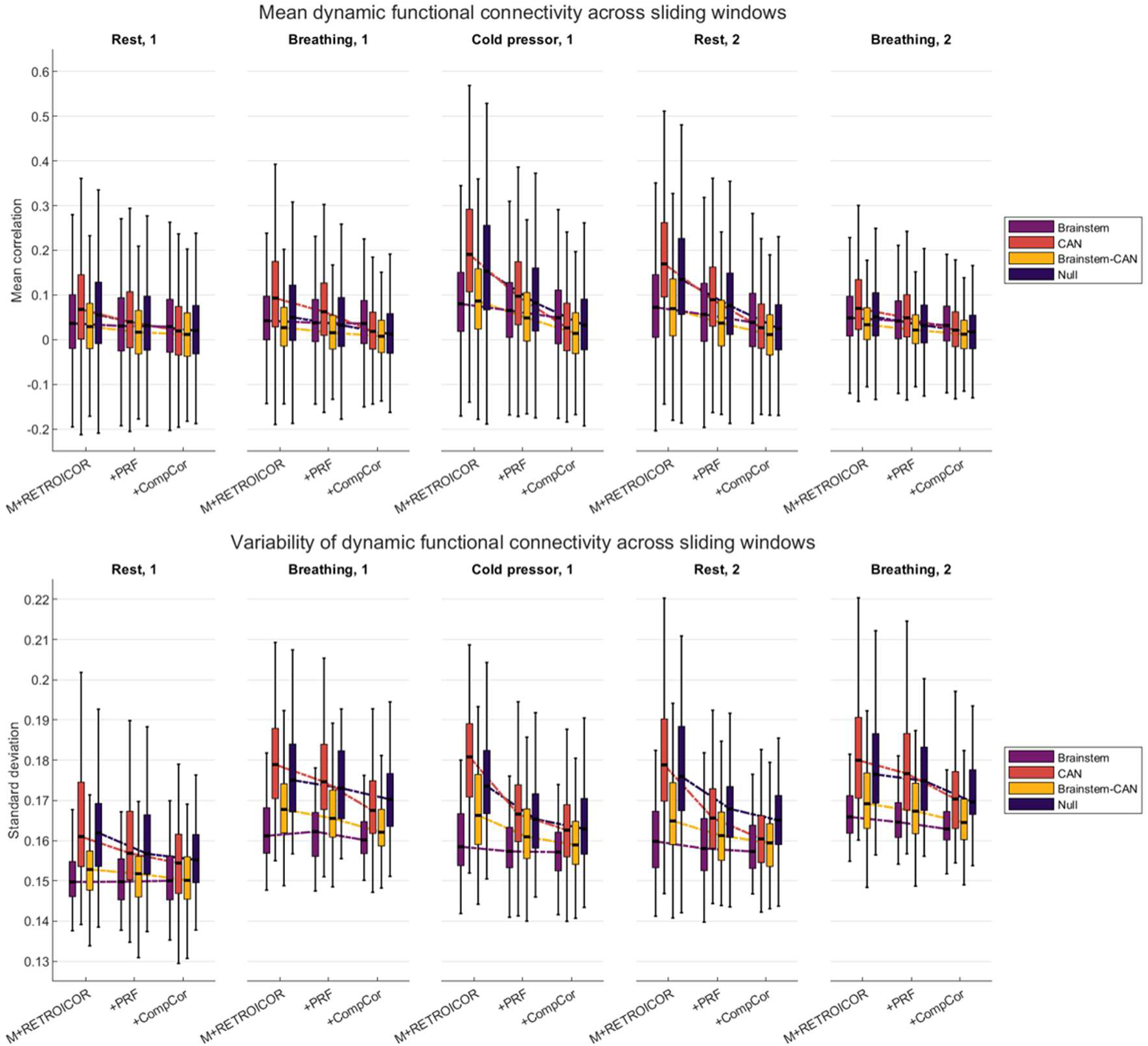
sLFO denoising reduces variability in mean FC and overall variability in FC values. Above: the distribution of FC averaged across all sliding windows is shown for different ROI pair types. Below: the distribution of the standard deviation of FC across all sliding windows is shown for different ROI pair types.

#### Linear mixed-effects model outcomes

For each denoising pipeline, measure of HRV, scan, and pair of ROIs (i.e. connectivity edge) we obtained beta values reflecting the modulation of dFC by HRV_trait_ (reflecting the baseline autonomic tone of each participant during each scan) and HRV_state_ (reflecting dynamic changes in autonomic outflow throughout the scan). The ROIs of dFC edges which yielded significant beta values for HRV_trait_ are reported in Supplementary Materials 10. These values reflect inter-subject changes in dFC linked to subjects’ mean autonomic activity throughout each scan. Notably, within the same measures and denoising pipelines, no significant results were reproduced within the same pair of regions for scans of the same type (i.e. rest during sessions 1 & 2 or breathing during sessions 1 & 2). However, several results were reproduced across pipelines or across the different HRV measures within the same pipeline (Figure 9A); of these, M+RETROICOR and PRF denoising were more similar than the CompCor pipeline. As well, the CompCor pipeline led to fewer significant beta coefficients for the HF-HRV and IBI measures than the M+RETROICOR and PRF pipelines. Some scan-specific results emerge: a positive relationship between HRV_trait_ and FC was found only during breathing task performance; otherwise, higher measures of HRV_trait_ was linked to lower FC across all pipelines.

**Figure 9:**
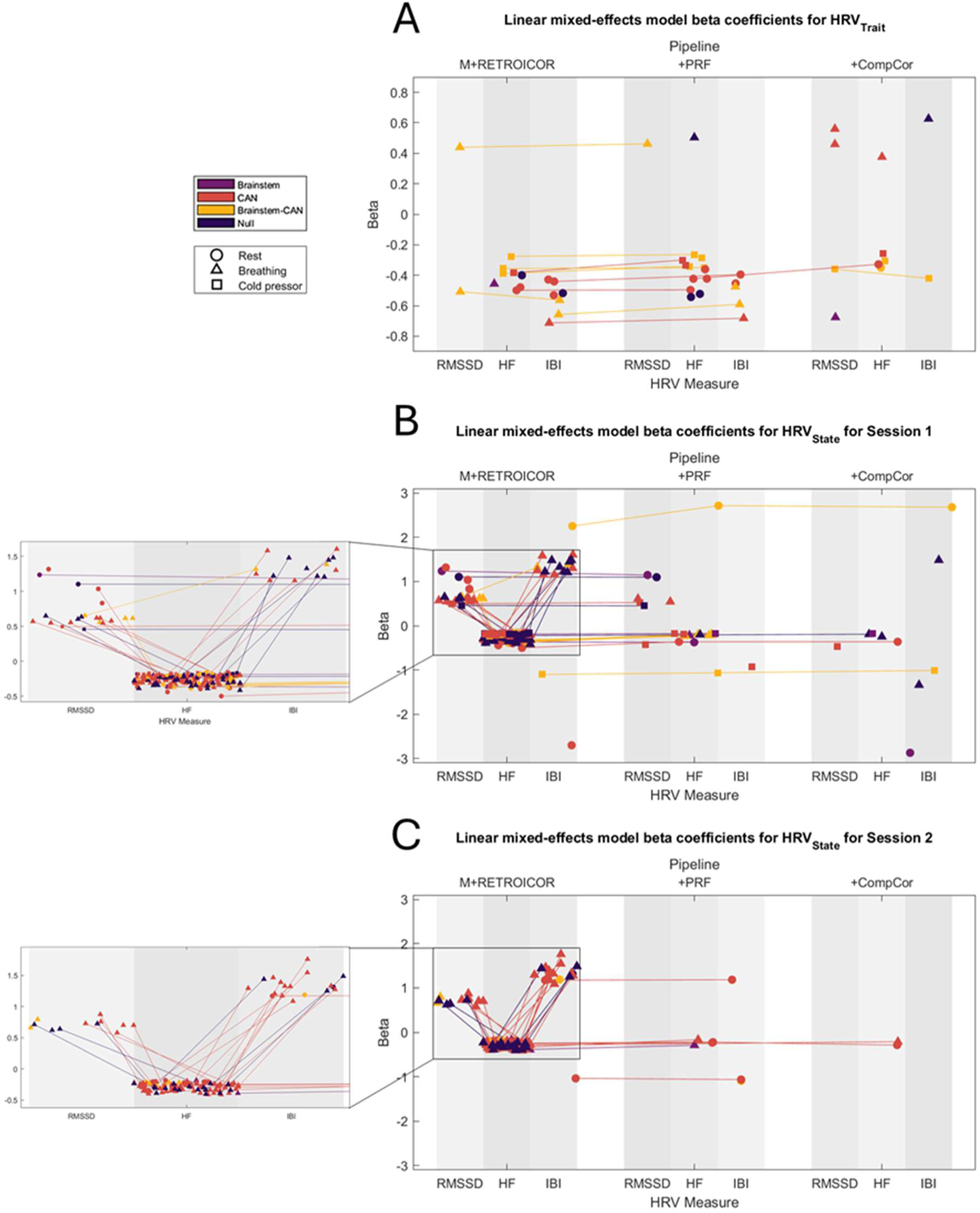
While RMSSD, HF-, and IBI measures of HRV_trait_ typically modulate autonomic-associated FC negatively, RMSSD and IBI measures of HRV_state_ modulate autonomic-associated FC positively; sensitivity to dFC modulation by HRV_state_ is significantly reduced by sLFO denoising. Model coefficients are shown for HRV_trait_ (A, p < 0.05, corrected for multiple comparisons) and HRV_state_ (B: session 1 and C: session 2, p ≤ 0.0005, not corrected for multiple comparisons) for each pipeline and HRV measure. Each marker represents a link between two ROIs for which connectivity was linearly modelled by a measure of HRV during a scan; marker shape reflects the scan’s task condition and marker color reflects the ROI pair type. Lines between markers reflect instances where the same pair of ROIs were implicated by multiple measures or multiple pipelines.

The results reported here for HRV_state_ model coefficients should be interpreted with caution, as we present results not corrected for multiple comparisons to better guide future study of subsets of the ROIs included here. Due to the large number of FC edges examined in the present study (561 in total, including null ROI connections), correcting for multiple comparisons would likely obscure results of interest in smaller, more targeted studies of the CAN and physiological brainstem nuclei. Further, we note that the highest level of significance which could be achieved using our null statistical model for HRV_state_ is p < 0.0005, which cannot be meaningfully Bonferroni-corrected in this implementation.

Results are presented separately for session 1 (Figure 9B) and session 2 (Figure 9C) to better assess the reproducibility of HRV_state_ model coefficients; however, at p **≤** 0.0005, the same dFC edges were reproduced within scans of the same type only for the M+RETROICOR pipeline, typically within the breathing pipeline (88 dFC edges reproduced; 86 for HF-HRV and one each for RMSSD and IBI). Of these, 15 dFC edges involving a null ROI were reproduced, including both brainstem and CAN connectivity to null ROI 3 (a sphere positioned in the primary motor cortex). Only one dFC edge was reproduced across resting-state scans, indicating dFC between the thalamus and supramarginal gyrus (CAN-associated ROIs 6 & 7) was negatively modulated by HF-HRV in both sessions. The intraclass correlation coefficient (ICC) across these reproduced edges was 0.92, indicating model coefficients were reproduced with high reliability across sessions; an ICC value of 0.92 was also obtained for the 405 M+RETROICOR edges reproduced at p < 0.05. Despite the general lack of ROI-specific reproducibility across scans for sLFO pipelines, similar trends in the effect of different HRV_state_ measures on dFC can be observed in both sessions. Both RMSSD and IBI measures of HRV_state_ were typically positively correlated with dFC (particularly for the M+RETROICOR pipeline), while HF-HRV was negatively correlated with dFC.

During session 1, several HRV_state_ coefficients were found to be reproducible across all three pipelines at p **≤** 0.0005. At rest, dFC between the right locus coeruleus (brainstem ROI 3) and secondary somatosensory cortex, posterior insula, and putamen (CAN-associated ROI 8) was positively modulated by IBI; dFC between the left ventral tegmental area (brainstem ROI 4) and the primary motor cortex and temporal pole (CAN-associated ROI 15) was negatively modulated by IBI during the cold pressor task.

To further assess the effect of denoising on ROI sensitivity to HRV modulation of dFC, as well as the effect of thresholding for statistical significance, a ranking of ROIs by node strength across all scans is shown for each HRV measure, pipeline, and several significance thresholds in Figure 10. For RMSSD and HF measures of HRV, CompCor denoising most aggressively reduced sensitivity to HRV modulation of dFC; only one dFC edge for each measure had a nonzero ranking at the highest significance level. For RMSSD, CompCor implicated HRV in modulation of FC between regions near the cingulate cortex (CAN-associated ROI 1) and amygdala (CAN-associated ROI 12), while for HF-HRV, dFC between the anterior insula and ventrolateral prefrontal cortex (CAN-associated ROI 5) and null ROI 3 (a sphere positioned in the primary motor cortex) was most significant. In contrast, multiple dFC edges for PRF and M+RETROICOR denoising were significantly modulated by these two HRV measures at p < 0.0005. The node strength rankings across pipelines and measures did not consistently differentiate between different ROI types: brainstem nuclei, parasympathetic and sympathetic CAN-associated ROIs, and null ROIs are all implicated in HRV modulation of dFC at p < 0.0005. We further note that sLFO denoising did not only reduce the node strength of regions implicated in the M+RETROICOR pipeline: PRF and CompCor denoising resulted in a more significant connection with the right locus coeruleus (brainstem ROI 3) for IBI.

**Figure 10:**
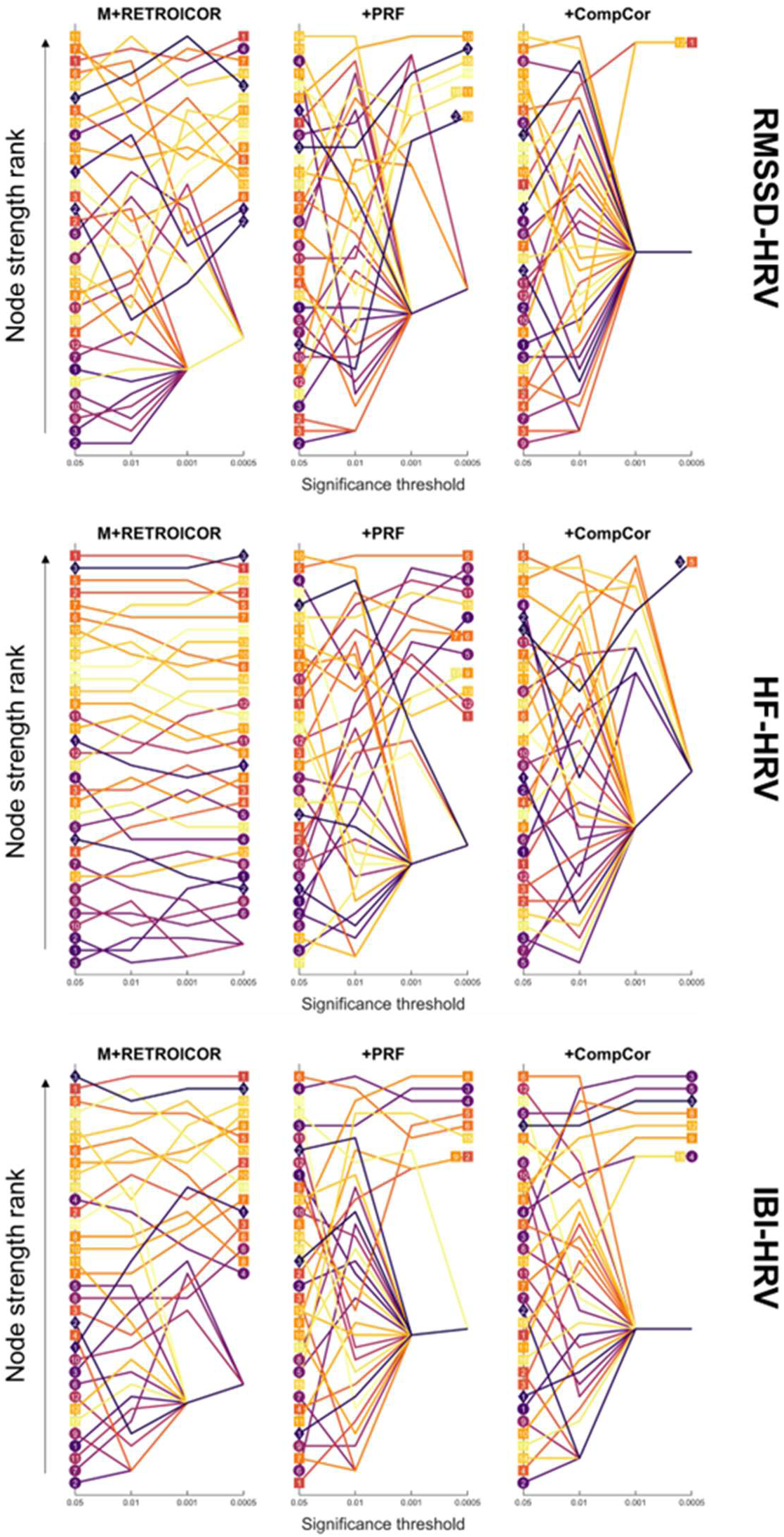
Node strengths of autonomic-associated regions are altered by denoising pipeline and vary significantly between measures of HRV. For each pipeline and HRV measure, node strengths were summed across all scans and then ranked across ROIs. ROI rankings are depicted at each significant level with markers corresponding to ROI type (brainstem ROIs: square; CAN ROIs: circle; null ROIs: diamond; all ROI numbers are defined in Supplementary Materials 1-4). At the highest significance level (p < 0.0005, not corrected for multiple comparisons), only ROIs with nonzero node strength are shown.

Finally, to assess the similarity of HRV_state_ model coefficients across scans and pipelines, the Spearman rank correlation coefficients for the node strength of all ROIs is shown in Figure 11 (FC matrices were thresholded to p < 0.01 prior to node strength calculation). In general, the Spearman coefficient is higher within individual scans (across different pipelines), than between different pairs of scans or across types. One exception to this is the breathing task, where node strength was ranked highly similarly within the M+RETROICOR pipeline for both sessions, for all HRV measures. However, in general, low similarity (< 0.4) was observed when comparing different scans. Within the same scan, the similarity between pairs of pipelines was not always consistent: M+RETROICOR and CompCor denoising were often least similar to each other but were more similar to each other than to PRF denoising during session 2’s breathing task for RMSSD HRV and IBI models. At rest, across all measures, as suggested by the node strength rankings depicted in Figure 10, PRF denoising typically offered a middle ground between the M+RETROICOR and CompCor pipelines.

**Figure 11:**
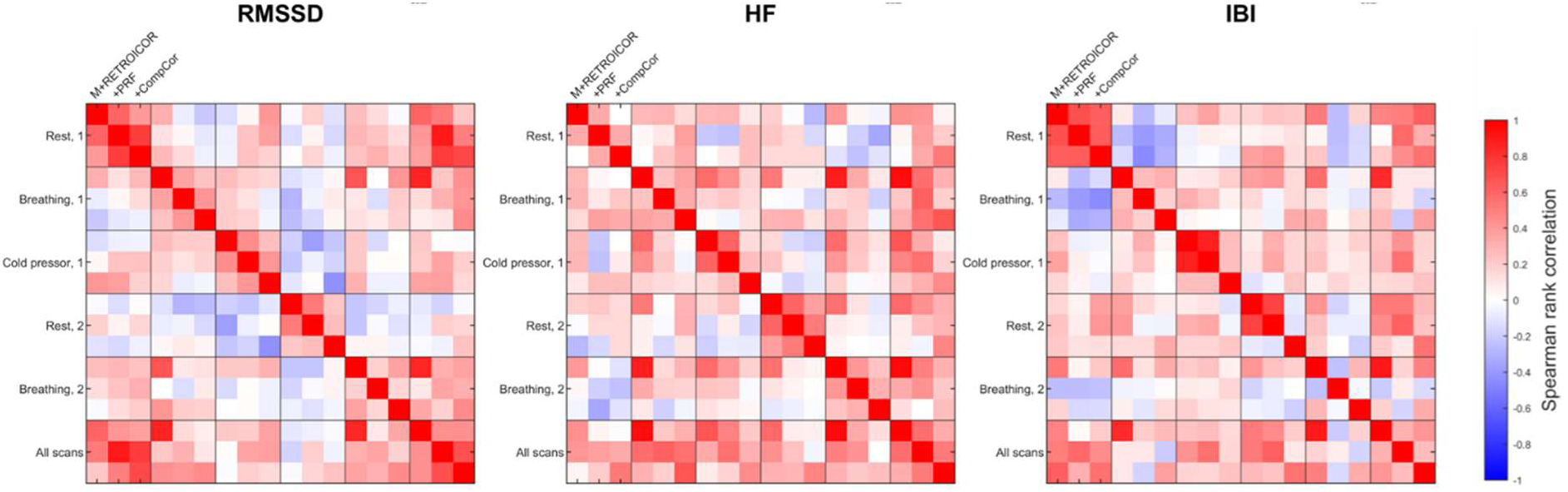
Node strength outcomes are typically more reproducible across pipelines within scans than within pipelines across scans. The Spearman rank correlation for the node strength of all ROIs is shown for pairs of scans and pipelines at the same significance level (p < 0.01, not corrected for multiple comparisons); intra-scan results may be compared within squares along the diagonal.

## DISCUSSION

### Summary of main findings

In the present investigation, we used a multimodal dataset including fMRI, EEG-fMRI and concurrent physiological recordings during rest and autonomic-related tasks to demonstrate the effects of sLFO denoising on three parallel characterizations of BOLD dynamics within and beyond the CAN. First, we found that PRF and CompCor denoising similarly reduce the coherence of BOLD signal in autonomic regions with changes in HR and RF (Figure 2, Figure 3). However, we also observed that sLFO denoising increased coherence with HR and RF when averaged across autonomic regions (Figure 2B). Moreover, we found that coherence with HR and RF was similar across autonomic-associated and non-autonomic-associated regions in most frequency bands (Figure 4), showing that the observed coherence patterns are not specific to autonomic regions only. Lastly, we showed that during coherence with RF and HR, vigilance tended to be lower and motion tended to increase, with some differences across regions, frequency bands, and denoising pipelines (Figure 5). In sum, our results suggest that BOLD coherence with physiological signals more likely reflects a combination of non-neuronal physiological artefacts and global processes related to arousal than local neuronal activity reflecting information exchange with autonomic systems, which would be specific to autonomic regions of the brain.

Next, we extended these results to CAN-related dFC by examining the relation of co-activation patterns occurring within autonomic-associated regions to fluctuations in physiological state and motion. Across various dimensionalities of data decomposition, we showed that CompCor denoising was associated with reduced sensitivity to changes in vigilance and motion, while PRF denoising reduced sensitivity to changes in HR coupled to changes in breathing (Figure 6). We further showed that co-activation patterns typically isolated an inverse relationship between vigilance and other physiological processes (HR, breathing, and motion, as shown in Figure 7), although PRF denoising and higher dimensionalities of data decomposition reduced this coincidence in physiological changes; higher BOLD activity in the cingulate cortex, supramarginal gyrus and somatosensory cortex, and ventrolateral prefrontal cortex was particularly associated with these observed differences in physiological state.

Finally, we used different measures of HRV to model changes in sliding window connectivity between autonomic-associated regions of the brain modulated by autonomic arousal. We found that while HRV_trait_ measures negatively modulated autonomic-associated dFC negatively, measures of HRV_state_ modulated autonomic-associated FC both positively (RMSSD and IBI) and negatively (HF-HRV). Further, denoising with sLFO regressors substantially reduced the significance of the modulation of dFC by HRV; this reduction in significance was more prominent in HRV_state_ than HRV_trait_ (Figure 9). For specific connections between autonomic-associated regions, HRV_state_ measures modelled changes in dFC with low reproducibility between different resting-state acquisitions, while high reproducibility (ICC = 0.92) was obtained with the M+RETROICOR pipeline during breathing task performance. While the PRF and CompCor denoising pipelines both reduced the significance of the HRV_state_ LME model coefficients, PRF-denoised outcomes were typically more similar to M+RETROICOR-denoised outcomes (Figure 11). Finally, we showed that the sensitivity of autonomic-associated regions’ dFC changes associated with HRV varied significantly between different denoising pipelines and measures of HRV_state_ (Figure 10), with notable instances of non-autonomic-associated regions exhibiting dFC modulated by HRV_state_.

Overall, our results highlight the susceptibility of dynamic signatures of autonomic processes using BOLD to methodological variability at the level of denoising. Moreover, the parallel methods of characterizing dynamic changes associated with changes in physiological state which we employed converge on evidence that these dynamics are not limited to autonomic-associated regions alone, but stem from brain-wide processes associated with both artefacts and arousal levels.

### Evidence for global fMRI dynamics coupled to arousal

Studies of the CAN have long pointed towards heterogenous regional involvements associated with parasympathetic and sympathetic processes which extends to the recruitment of sensorimotor, premotor, and temporal regions during task-related periods of heightened arousal (Beissner et al., 2013; Chouchou et al., 2025). However, it is useful to make a distinction between regionally- and mechanistically-specific neuronal activations directly related to autonomic processing (e.g. activations of brainstem nuclei, bilateral information transfer between the amygdala and hypothalamus, or the amygdala’s couplings with regions of the cortex linked to emotion) and the large-scale effects of arousal on brain dynamics, including both the participation of other regional architectures such as the default mode network and salience network (Unsworth and Robison, 2017; Young et al., 2017) and the global fMRI signal (Bolt et al., 2025), in discussions of the central autonomic network and the neural correlates of arousal.

First, the coherence with physiological signals we observed appears to relate both to sLFO-associated physiological noise and arousal levels. The sLFO-denoising pipelines similarly reduced coherence with HR and RF (Figure 2, Figure 3) but did not wholly eliminate it; coherence in all three pipelines was associated with changes in vigilance (Figure 5). In the co-activation pattern and dFC analysis of the CAN regions we identified, we did not observe a differentiation between regions associated with parasympathetic and sympathetic subnetworks which might be expected for changes in brain state associated with autonomic processing. Rather, the CAP states which were associated with changes in physiological state showed primarily changes in cortical regions (Figure 7), which could be potentially consistent with changes in broad networks during arousal; however, the interpretation of this analysis is limited without the inclusion of non-autonomic brain regions in the CAP decomposition.

Furthermore, connections whose dFC was modulated by HRV exhibited low specificity to key CAN regions and poor reproducibility across scans. The reliability of these dFC results is itself suggestive that connectivity between the examined regions was mediated by arousal. Previous studies have characterized dFC as less reliable compared to static FC (Lurie et al., 2020); in particular, when dFC was assessed between pairs of regions in sliding windows during rest, the magnitude of dynamic changes in FC was strongly correlated to lower test-retest reliability (Zhang et al., 2018). At the same time, the test-retest reliability of static FC has been shown to improve when excluding periods of sleepiness from resting-state data (Wang et al., 2017). Crucially, this result was demonstrated with periods of sleepiness identified by windows of time with higher RMSSD HRV, which did not correspond to differences in head motion across subjects. Additionally, sleepiness in Wang et al.’s study increased as scans went on. This same phenomenon of a decrease in arousal over time during fMRI scanning has previously been associated with artefactual inflation of FC caused by sLFO artefacts (Korponay et al., 2024).

Overall, the BOLD fluctuations in autonomic-associated brain regions which we observed across denoising pipelines seem less specific to the CAN itself and more reflective of global brain dynamics coupled to arousal. Therefore, future studies of the CAN may benefit from a stronger autonomic stimulus to induce a stronger neural response. Furthermore, future studies of the CAN should strongly consider study designs which incorporate multivariate analyses and include ROIs not expected to participate in autonomic-specific pathways to assess the specificity of their findings.

### Noise and signal in autonomic brainstem imaging

The inclusion of brainstem regions in the present study presented a unique opportunity to assess their suitability for detecting meaningful neuronal activations alongside cortical and subcortical CAN-associated regions. It is reasonable to expect strong physiological noise to obscure neuronal activity in brainstem nuclei (Beissner, 2015), but recent development of the Brainstem Navigator toolkit has shown that the connectivity of brainstem nuclei may be mapped even at 3T (Cauzzo et al., 2022; Hansen et al., 2024). Employing denoising strategies for autonomic brainstem nuclei may facilitate the detection of meaningful signals: the use of physiological noise correction has previously been shown to improve fMRI SNR in the locus coeruleus (Schumann et al., 2018). Moreover, dynamic analyses between brainstem nuclei and cortical regions have previously distinguished separable states relevant to autonomic function. Specifically, in a hierarchical clustering analysis performed on sliding-window dFC extracted from resting-state fMRI of patients with focal epilepsy and healthy controls, Mueller et al. identified two activity states in which the strength of connectivity between brainstem ROIs and the brain was significantly correlated with HRV. In the first – positively correlated with HRV – the strength of connectivity within gray matter and strength of connectivity within the CAN during this state together explained 80% of the overall variability in HRV for both study populations, while the second – negatively correlated with HRV – was linked to autonomic dysfunction and more prevalently observed in the focal epilepsy group. Notably, functional connectivity within cortical regions of the CAN did not correlate with HRV in the latter state, which was also marked by weaker connectivity between the brainstem and gray matter regions (Mueller et al., 2019).

In the present study, changes in BOLD in brainstem nuclei did not factor strongly in CAP states associated with physiological differences, suggesting these regions’ contributions to physiological state change was less observable than those of cortical regions, although this difference was somewhat lessened by the CompCor pipeline (Figure 7). Similarly, dFC variability was lowest in brainstem edges (Figure 8) while connectivity between and to brainstem regions was less commonly modulated significantly by HRV (Figure 10), which together suggests that our acquisition and preprocessing pipeline lacked sensitivity to neural autonomic processing in these regions. Future investigations of fMRI dynamics in autonomic nuclei may benefit from employing sequence parameters optimized for brainstem imaging (Matt et al., 2019; Turker et al., 2021).

### Suggested approaches to fMRI denoising in an autonomic context

The results of this study have important implications for both investigations of the neural correlates of autonomic activity and the interpretation of denoising in non-autonomic contexts. First, the impact of different preprocessing techniques is under-explored in fMRI literature of brain-body interactions, and the variation in autonomic-related dynamics across the sLFO-focused pipelines reported here should motivate further exploration of denoising effects in subsequent studies. In contexts where this methodological variability is not itself an effect of interest, employing several denoising pipelines may allow an assessment of this variability’s contribution to the desired target of study and improve the interpretation of findings in relation to physiological noise sources. At minimum, clearly reporting the rationale for selecting a specific denoising technique will aid researchers and reviewers in understanding the appropriateness of an fMRI workflow for a given research question. As previously discussed, we further suggest that researchers targeting the CAN should include regions not associated with autonomic regulation in their study design to better differentiate signal fluctuations associated with brain-wide arousal processes from local neuronal activations meaningfully involved in physiological regulation.

Second, the present study sheds light on several possible misconceptions related to the effects of physiological denoising in autonomic contexts. Although the use of any physiological denoising is sometimes eschewed to avoid removing sensitivity to signals of interest, all three pipelines in the present work incorporated physiological denoising and preserved substantial variability associated with physiological changes. Further, while sLFO-denoising reduced variability associated with physiological changes, its removal of physiological artefacts potentially increased sensitivity to neural signatures of autonomic arousal in several regions for which HRV was found to modulate dFC in only PRF and CompCor pipelines (Figure 10). However, the interpretation of such results is complicated by the findings of (Nalci et al., 2019), who reported that some FC estimates were correlated with the geometric norms of nuisance terms: significant results which arise only in the context of sLFO denoising may then result from either the uncovering of buried neuronal signals or from nuisance-related ‘injections’ into FC estimates. Regardless, the application of physiological denoising in autonomic contexts should be conceived in more nuanced terms than only a reduction of sensitivity. For those researchers who do not eschew physiological denoising, model-based techniques such as PRF denoising have sometimes been viewed as more aggressive than CompCor methods, since PRF regressors typically include a time series derived from HR and measures of autonomic arousal are likewise frequently derived from HR, albeit with some differences in temporal dynamics. However, we found that PRF denoising tended to be more conservative than CompCor, potentially by restricting the removal of physiological noise to faster (∼1-10s) temporal couplings with HR and RF and thereby preserving dynamics related to slower changes in global arousal.

Thirdly, we provide specific considerations for the implementation of sLFO-denoising techniques. Based on our examination of coherence in the case of the average signals extracted from all autonomic-associated regions, we caution against denoising sLFOs across ROIs without adjusting for regional effects such as hemodynamic delays (c.f. Frederick et al., 2012) when averaging BOLD signals across large regions or networks for later analysis. It seems likely that such an approach may introduce systemic physiological contributions to such averaged signals rather than removing them, potentially inflating the correlation between such signals themselves as well as their contamination by non-neuronal changes in physiological state. Further, to better preserve variability associated with changes in vigilance, we suggest using PRF denoising over CompCor denoising if concurrent physiological measurements are available to facilitate this. Importantly, this recommendation is consistent with previously reported reductions in whole-brain sensitivity to vigilance-related effects caused by CompCor denoising (Pourmotabbed et al., 2025).

Finally, we comment on the relevance of our findings for fMRI studies for which preserving the neural correlates of autonomic regulation are not of particular concern but for which characterizing physiological contributions to the BOLD signal remains pertinent. The application of physiological denoising is frequently discussed as analogous to removing variance associated with changes in arousal states, particularly when including sLFO regressors. The present study demonstrates that significant variance associated with arousal remains in fMRI dynamics after physiological denoising, including in those (e.g. motor) regions not strongly associated with autonomic processing. Our results therefore suggest that acquiring concurrent measures of physiology and including these as covariates in models of fMRI dynamics may improve the interpretability of analyses outside a purely autonomic context.

### Limitations and future directions

The present study examines only a subset of methods which may be relevant for investigations of the central autonomic network or other neural pathways related to physiological regulation. Within denoising methods, data-driven techniques such as ICA (Golestani and Chen, 2022), scrubbing of high motion fMRI volumes (Power et al., 2014), further interrogating the choice of RETROICOR model order, or comparing regionally-specific (Bartoň et al., 2019) or more aggressive (Kassinopoulos and Mitsis, 2019b) implementations of CompCor may equally be of interest when choosing an appropriate preprocessing pipeline. Beyond the plethora of alternative methods to assess dFC (Torabi et al., 2024), the set of ROIs selected as seeds within the CAN could be alternatively defined, or specific seeds associated with the CAN could be used to calculate voxelwise dFC across the brain (Chang et al., 2013). Furthermore, the use of different fMRI sequences and sequence parameters such as TR and encode direction can amplify, distort, or facilitate the modelling of physiological noise in ways that are relevant for future investigations (Constable et al., 2026; Todd et al., 2017). The treatment of physiological data and measures of HRV we employ are likewise investigated within a narrow context in the present study. Approaches to quality assurance for cardiac and respiratory measurements, peak detection, and calculation of HRV measures may introduce additional points of divergence from other studies. Further, the use of PPG to obtain HRV is known to differ from calculations using ECG data (Schumann et al., 2021b). Finally, multivariate methods such as partial least squares or canonical correlation analysis (Sui et al., 2012) could offer greater insight into the variation of distributed brain networks with diverse measures of physiological change.

We further note that the limited sample size of the present study could affect the sensitivity of our study design, particularly in uncovering regionally-specific autonomic signals, although we note that the difficulty of acquiring high-quality physiological signals in a multi-session study may result in similar sample sizes for other independent data acquisitions. However, as a result, the generalizability of these results should be approached with caution. This is especially true for applications of denoising in clinical or aging populations, given that age may mediate the effect of denoising on FC (Golestani and Chen, 2022), different populations may be differently susceptible to motion artefact (Weiler et al., 2022), and that differences in autonomic tone and function may be expected for a range of neurodegenerative and affective disorders (Cheng et al., 2022; Goffi et al., 2025; Liu et al., 2022). It has recently been shown that whole-brain patterns associated with sLFO processes may act as a biomarker of aging, with regions in the central autonomic network highly implicated in age prediction (Wang et al., 2026). Future studies in autonomically-dysregulated and/or aging populations would therefore particularly benefit by integrating an assessment of sLFO-denoising effects into their study design.

## CONCLUSIONS

In the present study, we explored the effect of sLFO denoising when dynamically characterizing physiological contributions to BOLD fluctuations in brainstem nuclei and brain regions associated with the CAN. We observed that while PRF and CompCor denoising similarly reduced BOLD coherence with peripheral physiological signals and each reduced the significance of HRV modulation of sliding-window dFC, these two methods of reducing sLFOs in fMRI data heterogeneously affected dynamics associated with autonomic arousal and head motion. Specifically, we found that CompCor denoising most reduced variability associated with fluctuations in vigilance and head motion, while PRF denoising outcomes were more similar to denoising with standard motion and RETROICOR regressors. This finding is particularly consequential for future investigations of the CAN, as CompCor denoising is sometimes seen in this context as more conservative alternative to model-based techniques for physiological denoising such as the use of PRFs. Moreover, our results implicated global arousal in BOLD dynamics of autonomic-associated regions rather than regionally-specific autonomic processing. Finally, low reproducibility of our models for HRV modulation of dFC across sessions point to a need for further replication of CAN dynamics, with particular attention to their relation to non-autonomic-associated brain regions. Our findings highlight the importance of assessing methodological variability introduced by denoising in fMRI studies of brain-body interactions to clarify the interpretation of neural correlates of autonomic regulation.

