## Supplementary Materials for "Denoising the central autonomic network: characterizing processes jointly associated with arousal and non-neuronal physiological artefacts in dynamic fMRI analyses"

### SUPPLEMENTAL MATERIALS

**Supplementary Materials 1: Brainstem nuclei included in the present work from the Brainstem Navigator toolkit based on their inclusion in autonomic or arousal processing (Bianciardi et al., 2016; Cauzzo et al., 2022; García-Gomar et al., 2022; Singh et al., 2022).**

| Brainstem regions | | |
|---|---|---|
| Legend number | Label | Association |
| 1 | Periaqueductal gray | A key component of the central autonomic network coordinating autonomic, pain-related, and motor responses (Benarroch, 1993; Coulombe et al., 2016) |
| 2 | Locus coeruleus (left) | With projections to multiple autonomic-associated regions along with somatosensory and motor nuclei (Benarroch, 1993), this nucleus has been implicated in respiratory control (Pattinson et al., 2009) and arousal (Munn et al., 2021) |
| 3 | Locus coeruleus (right) | |
| 4 | Ventral tegmental area – parabrachial pigmented nucleus complex (left) | Reported to have high functional connectivity with other brainstem nuclei included here along with cortical regions associated with the CAN; characterized as a hub between resting-state networks (Cauzzo et al., 2022) |
| 5 | Ventral tegmental area – parabrachial pigmented nucleus complex (right) | |
| 6 | Dorsal raphe | Considered part of the ascending arousal network (Bianciardi et al., 2016) and shown to exhibit a BOLD response during breath-hold tasks (Ciumas et al., 2023) |
| 7 | Lateral parabrachial nucleus (left) | Implicated in respiratory control (Benarroch, 2016) and in parasympathetic and sympathetic modulation of cardiac activity (Napadow et al., 2008; Valenza et al., 2017). |
| 8 | Lateral parabrachial nucleus (right) | |
| 9 | Medial parabrachial nucleus (left) | |
| 10 | Medial parabrachial nucleus (right) | |
| 11 | Viscero-sensory-motor nuclei complex (left) | Includes the nucleus of the solitary tract and vagus nerve nucleus (Singh et al., 2020), which are implicated in visceral sensory processing and parasympathetic outflow, respectively (Cutsforth-Gregory and Benarroch, 2017). |
| 12 | Viscero-sensory-motor nuclei complex (right) | |

**Supplementary Materials 2: Definition and labels of spherical seeds associated with the central autonomic network (CAN) which were included in the present work as 'CAN-associated regions'. These were adapted from a meta-analysis of brain areas previously associated with sympathetic and parasympathetic regulation (Beissner et al., 2013).**

| CAN-associated regions | | | | |
|---|---|---|---|---|
| Legend number | MNI coordinates | ROI radius (mm) | Label | Association |
| 1 | (0,10,40) | 5.94 | Midcingulate cortex, paracingulate cortex, supplementary motor area | Sympathetic regulation |
| 2 | (48, -26, 46) | 4.9 | Supramarginal gyrus, superior parietal lobule, primary somatosensory cortex | |
| 3 | (-20,-8,-12) | 4.02 | Amygdala, subgenual anterior cingulate cortex, nucleus accumbens, caudate, hippocampal formation | |
| 4 | (-2, 38, -18) | 3.91 | Ventromedial prefrontal cortex, pregenual/subgenual anterior cingulate cortex | |
| 5 | (44, 18, -6) | 3.76 | Anterior insula, ventrolateral prefrontal cortex | |
| 6 | (-4, -16, 6) | 3.74 | Thalamus (medial–dorsal nucleus), nucleus ruber, periaqueductal gray | |
| 7 | (-44, -36, 42) | 3.6 | Supramarginal gyrus, superior parietal lobule, primary somatosensory cortex | |
| 8 | (-32, -20, 14) | 3.51 | Secondary somatosensory cortex, posterior insula, putamen | |
| 9 | (-46, -66, -28) | 3.49 | Cerebellum (lobulus crus I) | |
| 10 | (20, 36, 34) | 3.44 | Dorsolateral prefrontal cortex | |
| 11 | (30,-22,-16) | 4.86 | Hippocampal formation | Parasympathetic regulation |
| 12 | (-20, -6, -18) | 4.29 | Amygdala, ventral tegmental area, hypothalamus | |
| 13 | (-40, 0, 12) | 4.18 | Anterior insula, caudate | |
| 14 | (-6, -44, 34) | 4.18 | Precuneus, dorsal posterior cingulate cortex | |
| 15 | (-56, 6, 8) | 4.13 | Primary motor cortex, temporal pole | |
| 16 | (50, -24, 2) | 3.8 | Medial temporal gyrus, superior temporal gyrus | |

| | | | |
|---|---|---|---|
| 17 | (44, -38, 14) | 3.79 | Supramarginal gyrus, angular gyrus |
| 18 | (-10, -62, -20) | 3.78 | Cerebellum (lobuli VI and vermis VI) |
| 19 | (40, 2, 12) | 3.71 | Anterior insula |

**Supplementary Materials 3: Definition and labels of regions included in the present work as regions not associated with autonomic regulation.**

| Null regions | | |
|---|---|---|
| Legend number | Label | Additional information |
| 1 | Inferior olivary nucleus (left) | Segmentation of this region obtained from the Brainstem Navigator toolkit (Bianciardi et al., 2015) |
| 2 | Inferior olivary nucleus (right) | |
| 3 | Primary motor cortex | Sphere of 5 mm radius centred at (-38, -24, 62), the central coordinate for activations in the left primary motor cortex during sensorimotor tasks (Hardwick et al., 2013) |

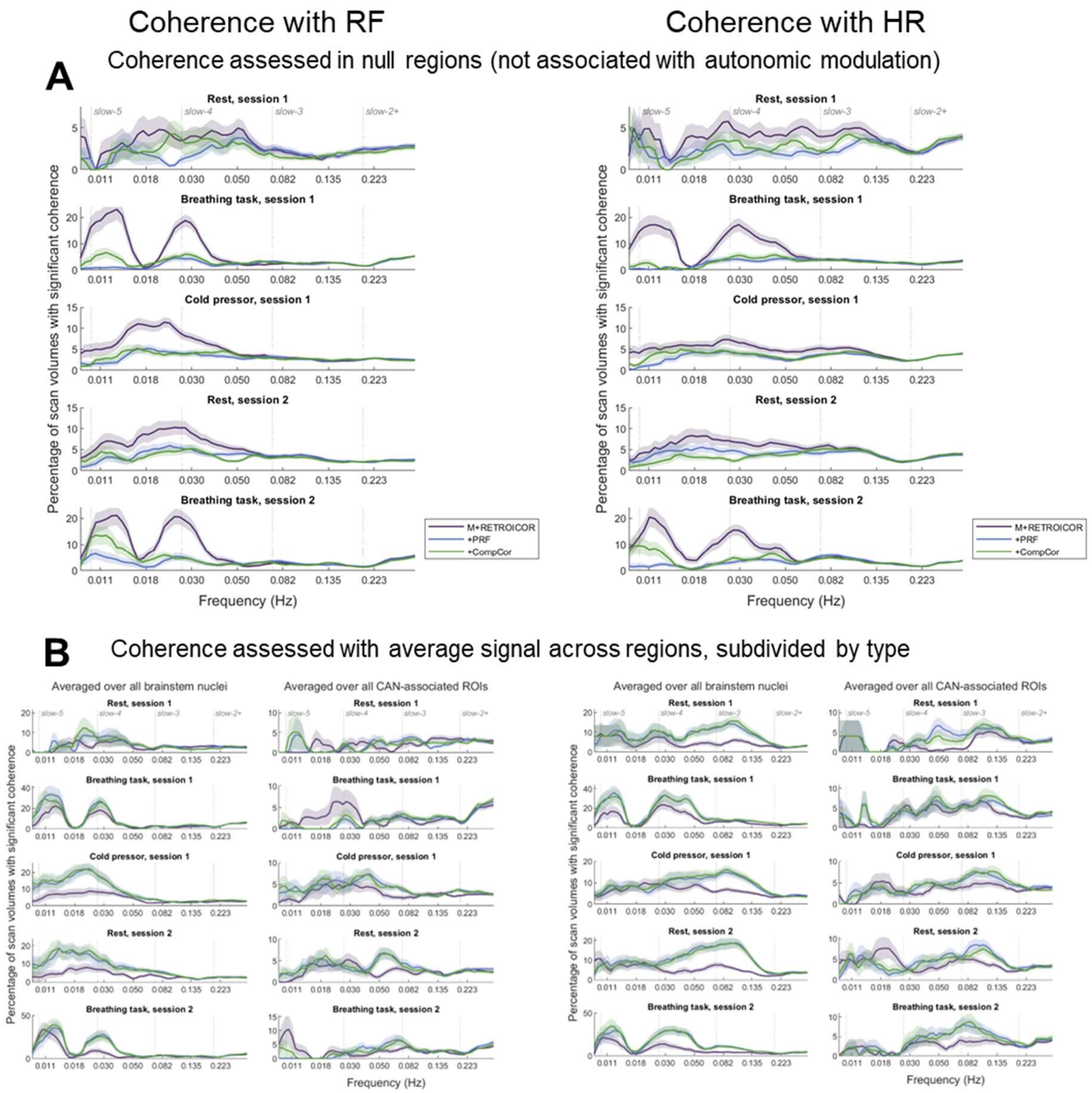

**Supplementary Materials 4:** The percentage of scan volumes which exhibited significant coherence is plotted against frequency for each scan and denoising pipeline. Results are shown for null regions (A) and for the average signal extracted from brainstem and autonomic-associated (brainstem and CAN) regions (B). Shaded regions represent the standard error for each pipeline and scan, over participants and ROIs (A) or over participants only (B).

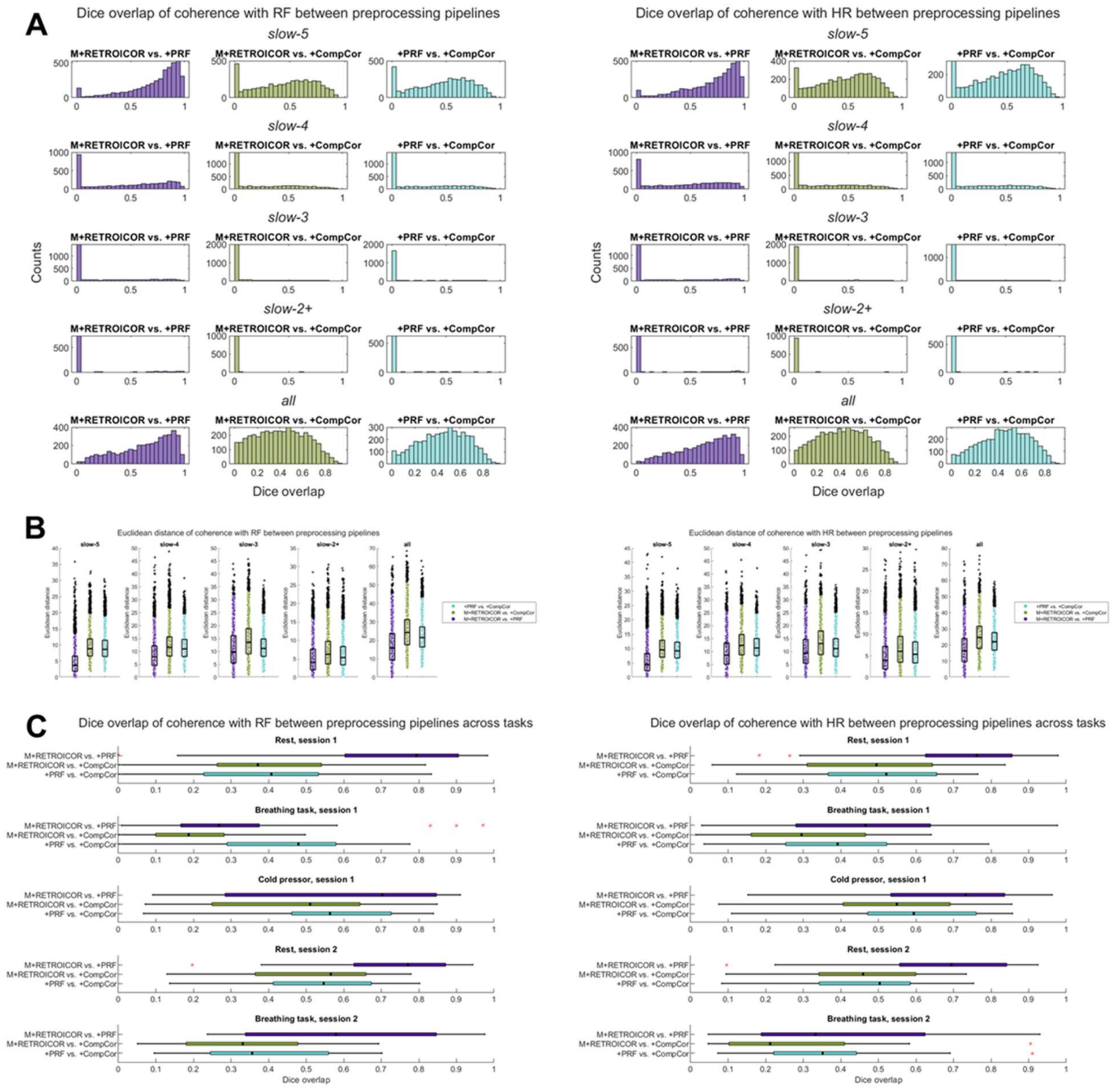

**Supplementary Materials 5:** Dice overlap scores were calculated for each individual scan's thresholded wavelet transform coherence patterns to reflect regions of significant coherence with RF (left) and HR (right). Dice overlap results are compared between pairs of denoising pipelines within frequency bands (A) and tasks (C). Additionally, the Euclidean distance between pipelines is plotted for the unthresholded wavelet transform coherence (outside the cone of influence) for each frequency band (B).

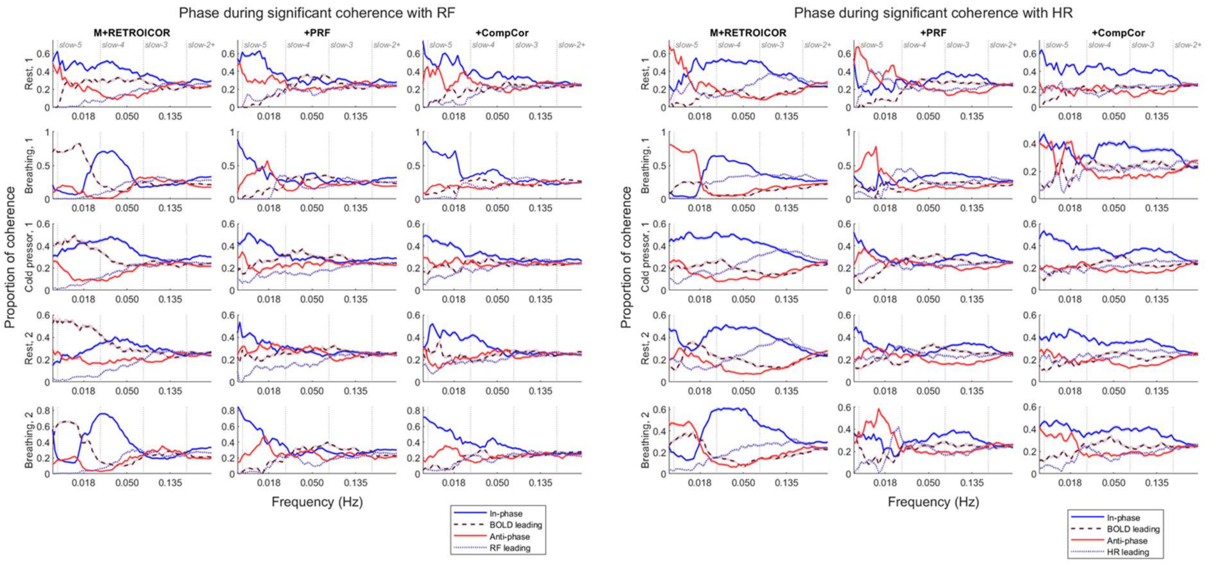

**Supplementary Materials 6:** The average proportion of significant coherence values found within each phase offset bin is plotted for each task and denoising pipeline for the cross-wavelet transform between RF (right) / HR (left) and signals extracted from each autonomic-associated ROI. Shaded regions represent the standard error for each pipeline and task over participants and ROIs.

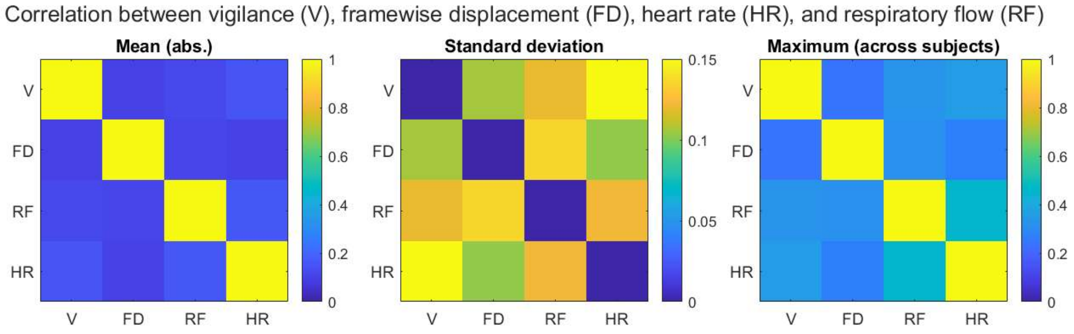

**Supplementary Materials 7:** Correlation between vigilance, framewise displacement, heart rate, and respiratory flow: mean absolute value of the correlation values averaged over all participants and session 2 scans (left), standard deviation (centre) and maximum absolute value (right).

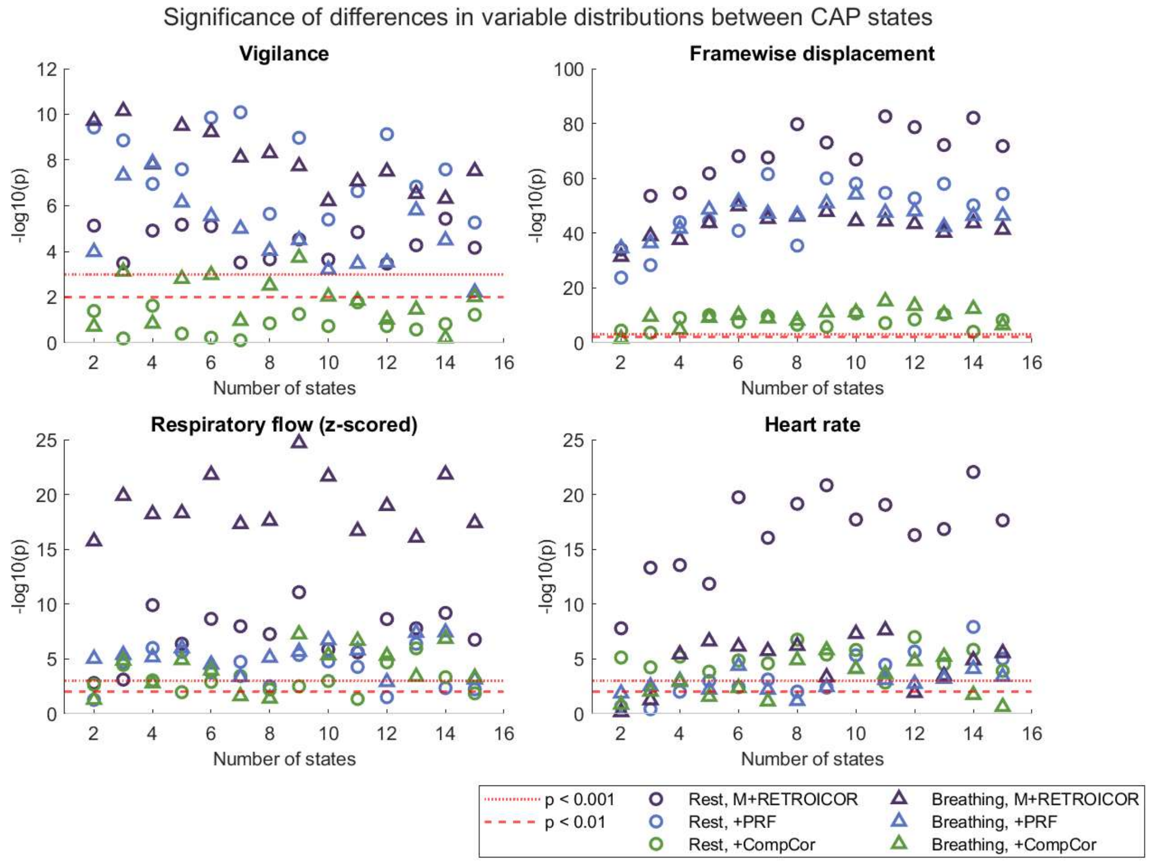

**Supplementary Materials 8:** The *p*-values from a one-way ANOVA are plotted here for each number of group-level CAP states ($n$ = 2…15) for physiological variables' distributions associated with CAP decompositions of rest and breathing scans acquired during session 2.

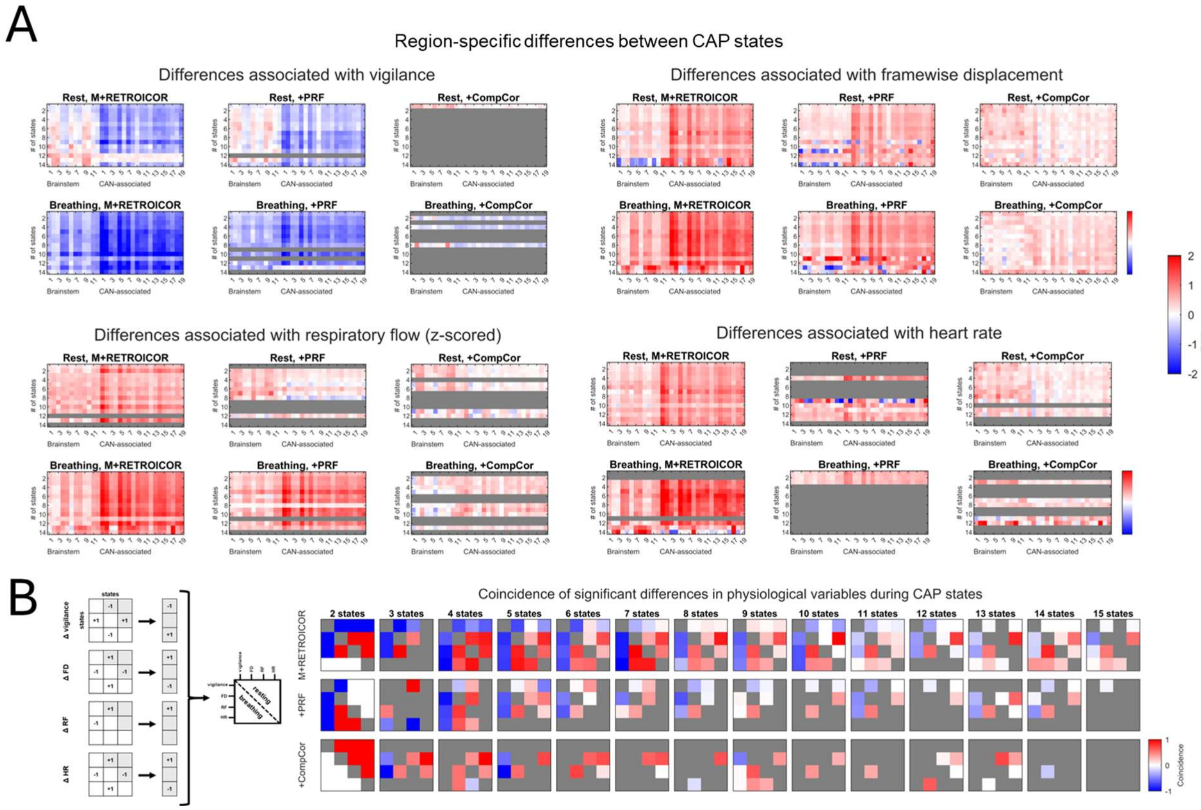

**Supplementary Materials 9:** Differences between centroids are plotted to show the change in each ROI only for pairs of CAP states exhibiting a change in physiological variable (A) while coincidence in changes between each pair of physiological variables is assessed across all CAP states (B). Gray regions represent instances where no significant differences in the distribution of a variable were observed across all CAP states.

**Supplementary Materials 10:** Significant linear mixed effects model coefficients ($p < 0.05$, corrected for multiple comparisons) and ROIs are reported for each measure of $HRV_{trait}$, pipeline, and scan.

| **Results between RMSSD HRV trait and autonomic-associated connectivity** | | | | | |
|---|---|---|---|---|---|
| *Pipeline* | *ROI 1* | *ROI 2* | *Scan* | *β* | *p-value* |
| M+RETROICOR | Brainstem 1 | CAN 13 | Breathing, 1 | -0.51 | 4.26E-05 |
| M+RETROICOR | Brainstem 12 | CAN 14 | Breathing, 2 | 0.44 | 2.99E-06 |
| PRF | Brainstem 12 | CAN 14 | Breathing, 2 | 0.46 | 1.97E-05 |
| CompCor | Brainstem 4 | Brainstem 12 | Breathing, 1 | -0.68 | 1.11E-07 |
| CompCor | CAN 3 | CAN 11 | Breathing, 1 | 0.46 | 7.03E-05 |
| CompCor | Brainstem 7 | CAN 5 | Cold pressor, 1 | -0.36 | 7.43E-06 |
| CompCor | CAN 9 | CAN 10 | Breathing, 2 | 0.56 | 1.94E-07 |
| **Results between HF-HRV trait and autonomic-associated connectivity** | | | | | |
| *Pipeline* | *ROI 1* | *ROI 2* | *Scan* | *β* | *p-value* |
| M+RETROICOR | Brainstem 4 | CAN 5 | Cold pressor, 1 | -0.39 | 3.24E-06 |
| M+RETROICOR | Brainstem 1 | CAN 8 | Cold pressor, 1 | -0.28 | 2.11E-05 |
| M+RETROICOR | CAN 3 | CAN 9 | Cold pressor, 1 | -0.38 | 1.15E-05 |
| M+RETROICOR | Brainstem 4 | CAN 19 | Cold pressor, 1 | -0.36 | 5.13E-06 |
| M+RETROICOR | CAN 2 | CAN 11 | Rest, 2 | -0.48 | 3.27E-08 |
| M+RETROICOR | CAN 7 | CAN 13 | Rest, 2 | -0.50 | 5.91E-05 |
| M+RETROICOR | CAN 11 | Null 3 | Rest, 2 | -0.40 | 4.68E-06 |
| M+RETROICOR | Brainstem 6 | Brainstem 9 | Breathing, 2 | -0.46 | 5.31E-05 |
| PRF | CAN 12 | Null 2 | Breathing, 1 | 0.50 | 6.29E-05 |

| | | | | | |
|---|---|---|---|---|---|
| PRF | Brainstem 4 | CAN 5 | Cold pressor, 1 | -0.34 | 2.77E-05 |
| PRF | Brainstem 1 | CAN 8 | Cold pressor, 1 | -0.26 | 1.86E-05 |
| PRF | Brainstem 2 | CAN 8 | Cold pressor, 1 | -0.29 | 1.75E-05 |
| PRF | CAN 3 | CAN 9 | Cold pressor, 1 | -0.30 | 4.97E-05 |
| PRF | CAN 2 | CAN 11 | Cold pressor, 1 | -0.33 | 5.38E-05 |
| PRF | Brainstem 4 | CAN 19 | Cold pressor, 1 | -0.35 | 1.94E-07 |
| PRF | CAN 2 | CAN 11 | Rest, 2 | -0.42 | 1.54E-08 |
| PRF | CAN 7 | CAN 13 | Rest, 2 | -0.50 | 5.59E-05 |
| PRF | CAN 12 | CAN 14 | Rest, 2 | -0.42 | 5.91E-05 |
| PRF | CAN 13 | CAN 16 | Rest, 2 | -0.36 | 8.15E-05 |
| PRF | CAN 2 | Null 3 | Rest, 2 | -0.54 | 1.17E-05 |
| PRF | CAN 3 | Null 3 | Rest, 2 | -0.52 | 3.84E-05 |
| CompCor | Brainstem 1 | CAN 8 | Cold pressor, 1 | -0.31 | 1.72E-05 |
| CompCor | CAN 10 | CAN 11 | Cold pressor, 1 | -0.26 | 1.61E-05 |
| CompCor | Brainstem 6 | CAN 10 | Rest, 2 | -0.35 | 1.18E-05 |
| CompCor | CAN 12 | CAN 14 | Rest, 2 | -0.33 | 2.37E-05 |
| CompCor | CAN 2 | CAN 3 | Breathing, 2 | 0.38 | 1.69E-05 |
| **Results between IBI trait and autonomic-associated connectivity** | | | | | |
| *Pipeline* | *ROI 1* | *ROI 2* | *Scan* | *β* | *p-value* |
| M+RETROICOR | Brainstem 1 | CAN 13 | Breathing, 1 | -0.56 | 1.50E-05 |
| M+RETROICOR | Brainstem 1 | CAN 15 | Breathing, 1 | -0.66 | 1.59E-05 |
| M+RETROICOR | CAN 5 | CAN 9 | Rest, 2 | -0.53 | 4.20E-06 |
| M+RETROICOR | CAN 2 | CAN 18 | Rest, 2 | -0.44 | 2.42E-05 |
| M+RETROICOR | CAN 7 | CAN 18 | Rest, 2 | -0.43 | 5.51E-05 |
| M+RETROICOR | CAN 18 | Null 3 | Rest, 2 | -0.52 | 5.32E-05 |
| M+RETROICOR | CAN 9 | CAN 19 | Breathing, 2 | -0.71 | 7.94E-05 |
| PRF | Brainstem 1 | CAN 15 | Breathing, 1 | -0.59 | 2.97E-05 |
| PRF | CAN 6 | CAN 12 | Rest, 2 | -0.45 | 7.48E-05 |
| PRF | CAN 2 | CAN 18 | Rest, 2 | -0.39 | 1.35E-05 |
| PRF | Brainstem 9 | CAN 13 | Breathing, 2 | -0.48 | 5.56E-05 |
| PRF | CAN 9 | CAN 19 | Breathing, 2 | -0.68 | 2.46E-05 |
| CompCor | Brainstem 7 | CAN 5 | Cold pressor, 1 | -0.42 | 1.12E-06 |
| CompCor | Brainstem 10 | Null 2 | Breathing, 2 | 0.63 | 4.93E-05 |

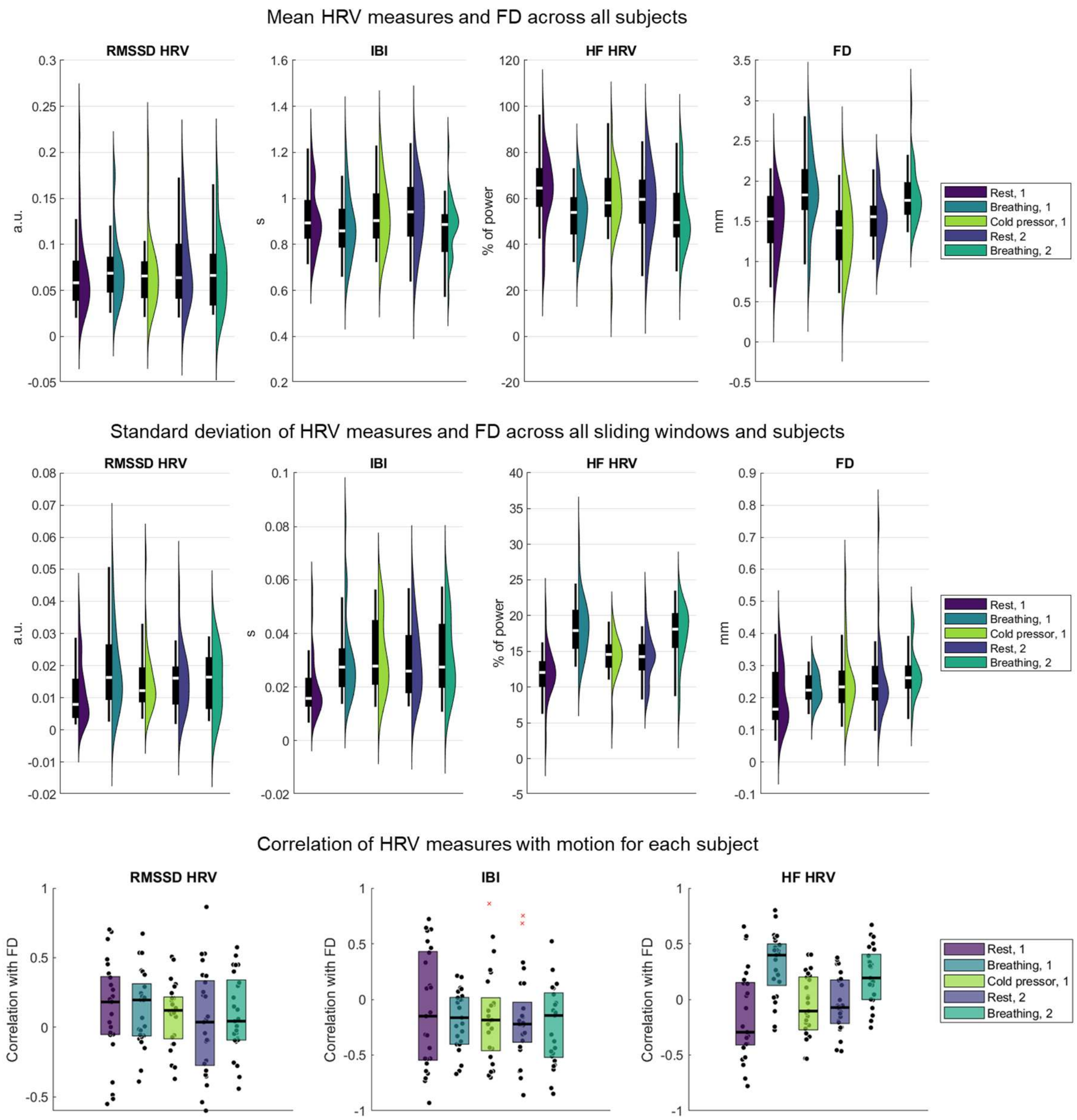

**Supplementary Materials 11:** Linear mixed-effect model input characteristics for each scan.
